# *Cryptococcus neoforman*s rapidly invades the murine brain by sequential breaching of airway and endothelial tissues barriers, followed by engulfment by microglia

**DOI:** 10.1101/2023.11.13.564824

**Authors:** Vanessa I. Francis, Corin Liddle, Emma Camacho, Madhura Kulkarni, Samuel R.S. Junior, Jamie A Harvey, Elizabeth R. Ballou, Darren D. Thomson, J. Marie Hardwick, Arturo Casadevall, Jonathan Witton, Carolina Coelho

**Affiliations:** MRC Centre for Medical Mycology at University of Exeter, University of Exeter, Exeter, EX4 4QD, UK; Faculty of Health and Life Sciences, University of Exeter, EX4 4QD, UK; Bioimaging Facility, University of Exeter, Exeter, EX4 4QD, UK; Johns Hopkins Bloomberg School of Public Health, Baltimore, Maryland, USA

## Abstract

The fungus *Cryptococcus neoformans* causes lethal meningitis in humans with weakened immune systems and is estimated to account for 10-15% of AIDS-associated deaths worldwide. There are major gaps in our understanding of how this environmental fungus evades the immune system and invades the mammalian brain before the onset of overt symptoms. To investigate the dynamics of *C. neoformans* tissue invasion, we mapped early fungal localisation and host cell interactions at early times in infected brain, lung, and upper airways using mouse models of systemic and airway infection. To enable this, we developed an *in situ* imaging pipeline capable of measuring large volumes of tissue while preserving anatomical and cellular information by combining thick tissue sections, tissue clarification, and confocal imaging. Made possible by these techniques, we confirm high fungal burden in mouse upper airway turbinates after nasal inoculation. Surprisingly, most yeasts in turbinates were titan cells, indicating this microenvironment enables titan cell formation with faster kinetics than reported in mouse lungs. Importantly, we observed one instance of fungal cells enmeshed in lamina propria of upper airways, suggesting penetration of airway mucosa as a possible route of tissue invasion and dissemination to the bloodstream. We extend previous literature positing bloodstream dissemination of *C. neoformans*, via imaging *C. neoformans* within blood vessels of mouse lungs and finding viable fungi in the bloodstream of mice a few days after intranasal infection, suggesting that bloodstream access can occur via lung alveoli. In a model of systemic cryptococcosis, we show that as early as 24 h post infection, majority of *C. neoformans* cells traversed the blood-brain barrier, and are engulfed or in close proximity to microglia. Our work establishes that *C. neoformans* can breach multiple tissue barriers within the first days of infection. This work presents a new method for investigating cryptococcal invasion mechanisms and demonstrates microglia as the primary cells responding to C. neoformans invasion.

## Introduction

*Cryptococcus neoformans* has been designated by the World Health Organisation as a critical priority pathogen that causes ∼112,000 deaths per year including 19% of HIV-associated deaths^1^. Healthy individuals acquire *C. neoformans* infections from environmental sources, and 56-70% of healthy children ages 1-10 years have serum antibodies against *C. neoformans* proteins^2^. Such early seropositivity suggests individuals frequently come into contact with *C. neoformans* and that infection in healthy individuals is either cleared or persists in latent, asymptomatic form^3,4^. Dissemination from airways requires *C. neoformans* to cross a series of tissue barriers to exit the airways, enter the bloodstream, and cross the blood-brain barrier where it causes meningoencephalitis^5^. Thus, *C. neoformans* can rapidly cross tissue barriers to invade the mammalian brain, suggesting the existence of sophisticated invasion mechanisms that are not understood.

We previously reported that viable *C. neoformans* could be recovered from mouse brains as early as 3 h, and fungal burden up to 7d after intranasal infection of mice^6^. Intravital microscopy studies detected *C. neoformans* traversal from the lumen of capillaries to the mouse brain parenchyma within a few hours after systemic infection^7–9^, consistent with *in vitro* models of endothelial tissue infection^10–13^.

How *C. neoformans* crosses tissue barriers on initial airway infection and to reside into the brain is still poorly defined. To efficiently identify the earliest sites of dissemination requires the capacity to observe and analyze rare, sparsely distributed invasion events. To achieve this goal, we implemented tissue clarification and decolorization, which remove lipids and certain pigments from tissues, resulting in a dramatic increase in tissue transparency which allows high-content imaging of thick samples^14^. This technique has great promise for studying host-pathogen interactions. This has been used to quantify *Aspergillus fumigatus* growth, and association with host immune cells in whole lungs^15^. This technique was also used to study cryptococcal infection-induced melanisation of *Galleria mellonella*, an insect model of cryptococcal infection^16^. Here, we combined tissue clarification with confocal microscopy to investigate early *C. neoformans* infection in mice airways and brain. In a model of intranasal infection, we present evidence for tissue barrier crossing in upper airways and lungs. We observed that within the first 24h after intravenous infection, majority of *C. neoformans* cells have traversed the blood-brain barrier (BBB) and are associated with brain-resident ionized calcium-binding adapter molecule 1 (Iba1^+^) macrophages. Our study is a key step towards defining the tissue routes and cellular interactions facilitating *C. neoformans* dissemination through mammalian hosts, and firmly implicating microglia as the primary brain immune cell responding to cryptococcal BBB traversal.

## Results

### High-content imaging of *C. neoformans*-host interactions in multiple infected tissues

To map early steps of host-invasion by *C. neoformans in vivo*, we combined mouse infections, tissue clarification, and high-content imaging (workflow illustrated in Fig. 1). We note that in most tissue clearing work perfusion is performed to reduce highly pigmented hemoglobin. We chose to not perform perfusion to avoid removal of *C. neoformans* yeast from the bloodstream. This was replaced with a decolourization step which degrades haemoglobin in the blood and other pigments to improve light penetration^17^. Following infection with *C. neoformans*, mice were sacrificed at 1-7 days, and the skull and lungs were collected and fixed. Skulls were decalcified, cut into thick sagittal tissue sections (10-13 per mouse), and clarified using X-CLARITY. These sections provided an optimal balance between preservation of anatomical context, efficiency of data collection, and sampling capacity. Lungs were also collected, clarified, decolorized and imaged whole, or in 2-3 coronal or axial sections. This was followed by staining for fungal cells with calcofluor white (CFW), for host cell nuclei with nuclear dyes which serves as anatomical landmarks, and, in some experiments, host immune cells by immunolabeling. In all, at least 200 μm depth of cleared skulls with 9 μm z-axial spacing per frame were analysed, corresponding to ∼4% of the brain volume (Fig. 1, see Methods). CFW dye can have background staining, including bone structures and debris of unknown nature. As a control, we imaged tissues from uninfected animals (Fig.S1). Fungal cells were detected and confirmed via manual verification of characteristic cryptococcal morphology (Fig.1 bottom panel). In summary, we developed a robust tissue clearing and imaging method that can be easily adopted to characterise pathogen invasion in rodent models.

**Figure 1.**
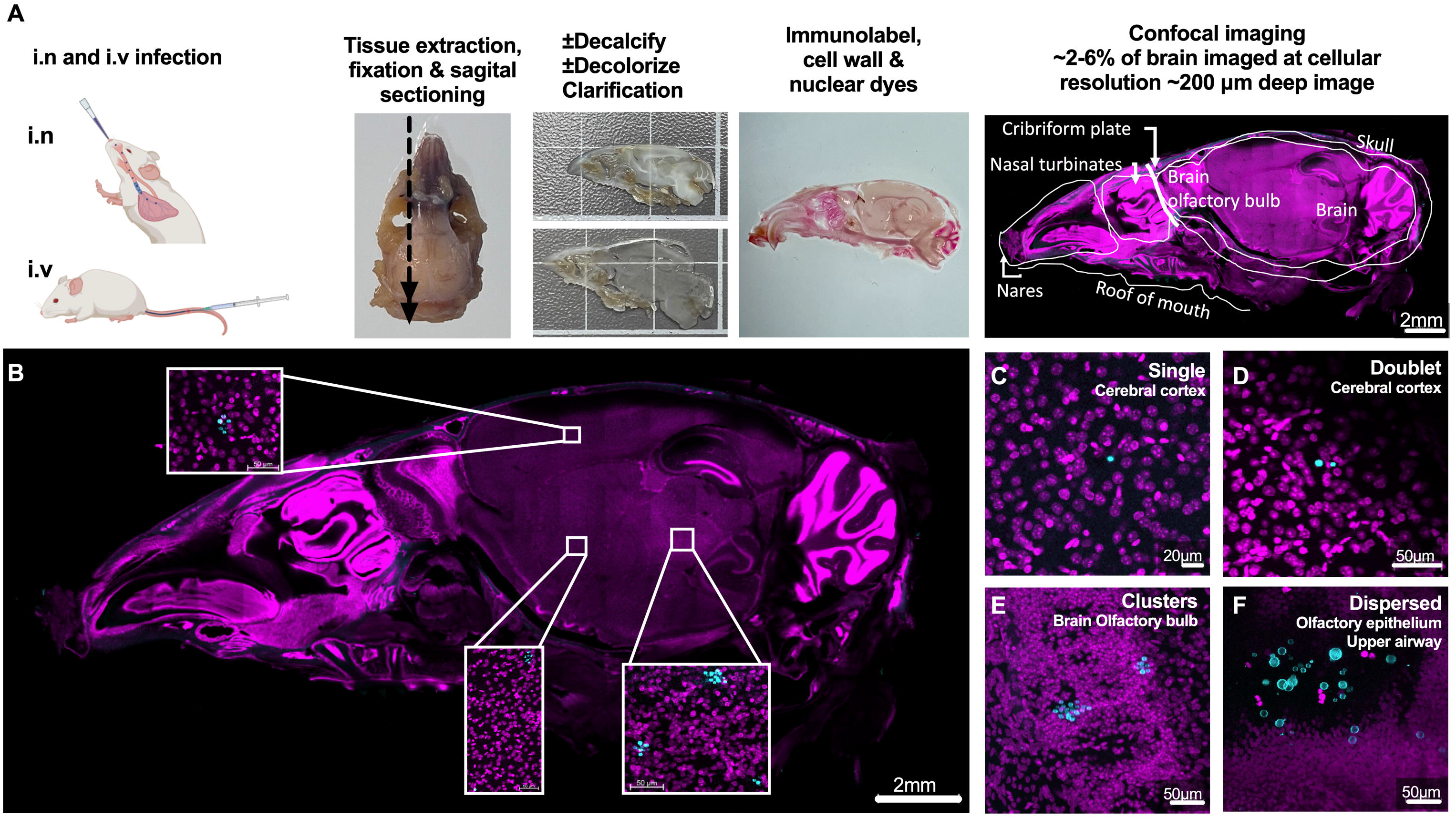
Workflow of high-content, subcellular resolution imaging of infected tissue slices. **A**) Schematic of experimental protocol. Representative images of **B)** a sagital cut slice of infected skull, showing insets of location of *C.neoformans;* and **C-F)** C. *neoformans* cells are observed in infected tissues as **C)** single, **D)** doublets, **E)** clusters, and **F)** dispersed. Images from skull of C57bl/6 male infected with 5×10^5^ H99E for 24h, via intravenous **(B-E)** or intranasal infection **(F).** Brain and skulls were cut in sagital sections, lungs into lobes or into coronal slices. Fungi identified by cell wall staining with CFW (50 µg/ml, cyan) or specific antibodies (see methods), tissues counterstained with different nuclear dyes (magenta), Pl, Helix NP™ Blue, and Helix NP™Green, depending on other fluorophores used. Maximum intensity projections with a depth of **C)** 36 µm (x5 z-steps), **D)** 63 µm (x8 z-steps), **E)** 105 µm (x36 z-steps) and **F)** 109.47 µm (x11 z-steps). Note third panel in A is same image as SFig.2. Scale bars in panels.

### Quantification of fungal titan cells *in situ*

Although *C. neoformans* titan cells (defined as titan cells (>10 μm diameter of cell body, determined by CFW) are important *in vivo*, they have not previously been characterized *in situ*, which was made possible with our new techniques. Volumetric imaging of fungal cells in mouse lung tissue (Fig.2A-C) revealed a wide range of cryptococcal cell sizes. To measure fungal size accurately, correction for loss of light intensity over the depth of tissues was performed (illustrated in Fig. 2D vs E). We measured fungal size in these images via the usual methods: manual diameter (by measuring cross-section, Fig.2F). We also used a semi-automated analysis, which are based on intensity thresholding fluorescent signals to define an object boundary, such as StarDist from ImageJ; area of the object can be used to calculate diameter of the object. We also decided to manually trace the circumference of fungi to directly compare to StarDist object boundary tracing (Fig.2G vs H), see also Methods and Supplemental results). Whilst cross-section and StarDist object boundary tracing show some discrepancy, manual object tracing supported the accuracy of the StarDist approach. Consistent with previous work, ∼40% of cryptococci, as determined by StarDist in the lung were titan cells (Fig.2I). We further confirmed our method was accurate, by testing conditions in which titan cells are rare (SFig.2 and Supplemental results). Thus, our imaging and analysis pipelones readily detect differences in fungal size *in situ* (see supplemental results for additional details). Our method can also be harnessed to obtain greater depth of imaging in widefield microscope compared to non-clarified samples. Using widefield fluorescence microscopy, we obtained z-stacks with high-quality images at >82 μm depth in a single tissue slice (see details in SFig.3, SVideos,and supplemental results).

**Fig 2.**
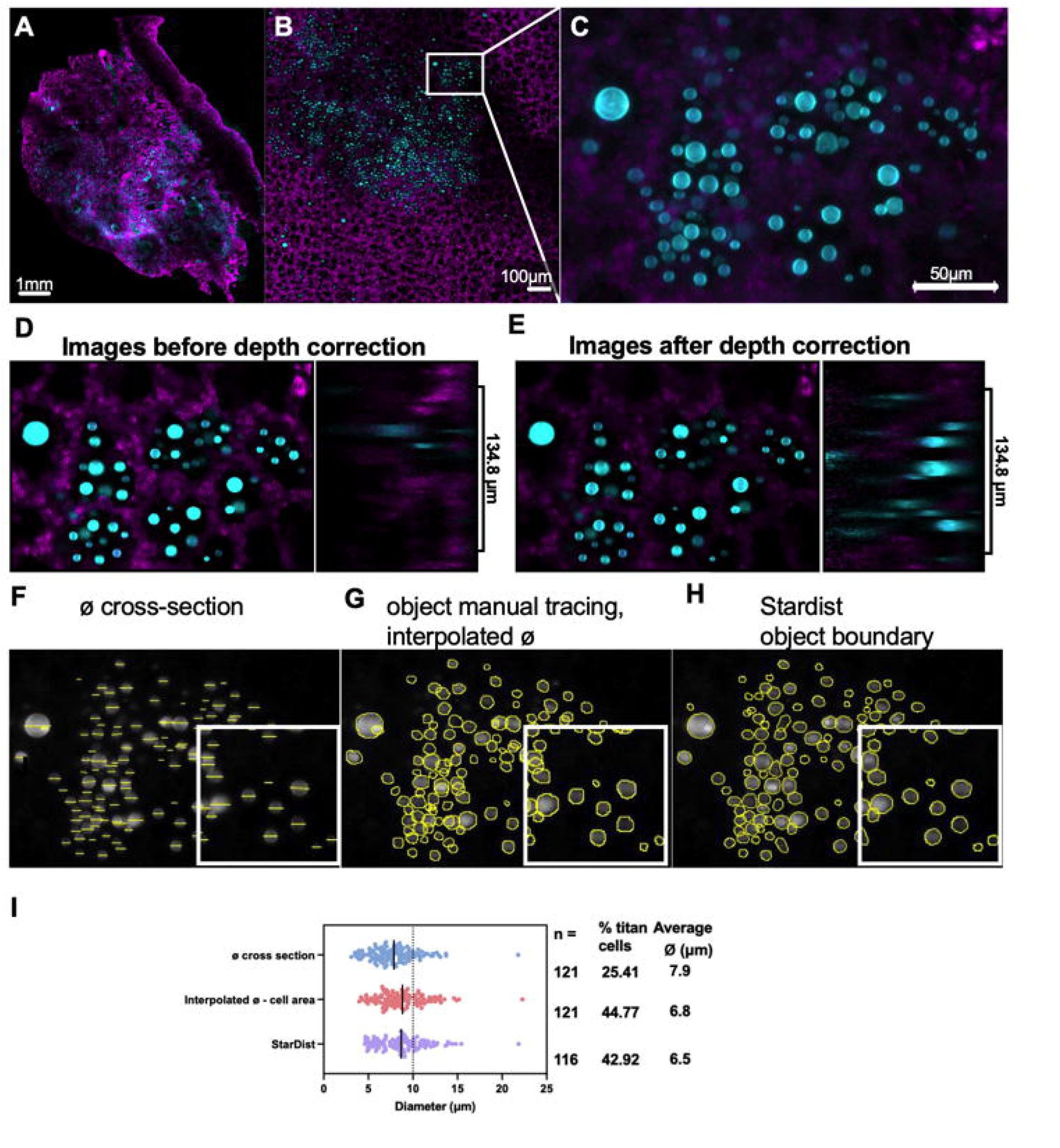
Pipeline to detect cryptococcal cell size distribution in infected tissues. Map of infected lung showing CFW stained cryptococcal cells and Pl stained nuclei of lung cells, with consecutive magnifications from tissue to subcellular resolution. Shown are several steps in image processing and analysis which allowed measurements of fungal celll morphology *in situ.* Different methods were compared, including the standard method of direct diameter, versus an object boundary methods which allow automated detection. Whe **A**) Single plane tiling map. **B)** Magnification and **C)**ROI used for quantification. **D)** Depth of tissue results in decreased signal intensity. Diminishing CFW signal in YZ projection (2.01µm x 68 z-step) in a single ROI (XY) single plane, andE) which is corrected by depth correction via a normalisation algorithm andimproves signal uniformity. Fungal size (diameter) was measured in this ROI via 3 methods: **F)**manually cross-section (0 cross section), **G)** manual object tracing, followed by area determination,and interpolation of diameter (interpolated 0), and **H)** automated analysis using lmageJ (NIH) StarDist macro, which uses thresholding to define object boundaries, followed by area and diameter calculation. **J)** Comparison of all three methods used to measure cryptococci size. n=total number of cells detected, cells >10 µmare dassified as titan cells ((dashed line). Images **(B-1)** from lung of C57bl/6J male mice, 5 dpiintranasal with 5×10^7^ C. *neoformans* ste50t.., a strain with virulence comparable to KN99 parental wild-type. Lungs stained with Pl (magenta) and CFW (cyan). Images correspond to **B-E)** extended depth of field image, covering 134.8 µm depth with 2.01 µm x 68 z-steps. Scale bar in images.

### Abundant *C. neoformans* titan cells present in upper airways by one day post infection

We previously showed that after intranasal inoculation, fungi are detectable in the upper airways as early as 24 hpi, with some fungal cells already forming titan cells^6^. For a more quantitative analysis, our high-content imaging approach was applied to skulls of mice infected via the intranasal route at 24 hpi and 7 dpi^6^. Imaging of sagittal slices taken from infected skulls, using the same dose and fungal strain as previous studies, showed cryptococci distributed throughout the upper airway turbinates at both 24 h and 7 d (Fig. 3 and Fig.4). At 24 hpi, yeast cells were adhered to the olfactory mucosa in nasal turbinates, including the superior turbinates (ethmoturbinates, most distal from nostrils – illustrated in Fig.3A and C), indicating capacity of yeast cells to overcome the first anatomical filtering barriers of the airways and to adhere to epithelial surfaces in these distal regions (Fig.3.a1). Remarkably, >50% of cryptococci in airways at 24 hpi were titan cells (Fig.3B), which indicates for the first-time that the environment in airway turbinates is a strong inducer of titan cell formation. We also surprisingly observed a large (≥10 μm) cryptococcal cell (Fig.3C) located in the lamina propria below the olfactory mucosa at 24 hpi (Fig.3C-panel c.3), suggesting invasion of airway mucosa, which had not been described before. At 7 dpi, we continued to observe abundant fungi in upper airways, including a high proportion of titan cells (Fig.4A-panel a1).

**Figure 3.**
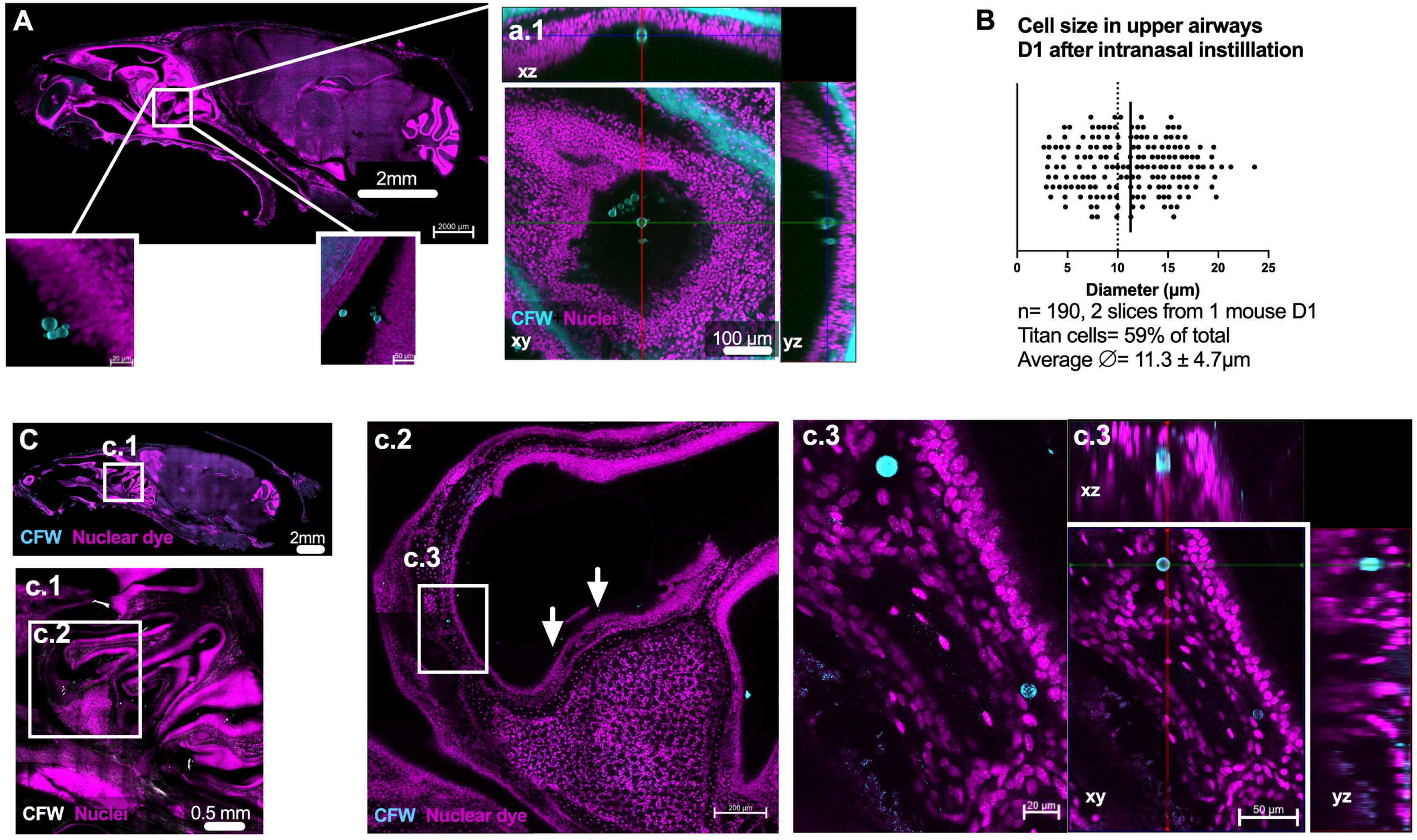
Presence of abundant titan cells in mouse airways 24 h after intranasal instillation of yeasts. C. *neoformans* and titan cells are abundant in upper airways within turbinates, closely aposed to, and invading lamina propria of the olfactory mucosa. A) Skull slice, showing several instances of cryptococcis apposed to mucosa and a1) ROI a magnified. [See supplementary video 1 to visualise YZ]. B) Titan cells are abundant in olfactory mucosa, as early as 24 hpi of H99E 5×10^5^ CFU. Cryptococci size measured using StarDist, combined data from two skull slices from the same mouse. C) Cryptococci can be observed throughout the upper airway, and invading mucosa, with c.1) highlighting location of fungal cells, and c.2) cryptococcii within turbinates (white arrows) and c.2-3) emeshed in lamina propria, below mucosal layer; cell body of diameter 12.98 µm (top) and 9.85 µm (bottom) measured by StarDist. A) Skull single plane a.1) xy single planes, followed by and xyz projection (right panel). B) Quantification of cryptococci size. 2 skull slices from the same animal were imaged with a depth of 225 or 208 µm, 26× 9µm or 20×10.95 z-step respectively. C) Skull slice (same animal as A, new slice, with maximum projection, 2×27 µm z-step). c.1) max projection 225 µm, 26×9 µm z­ step; c.2) 6 µm max projection (2×6 µm z-step); c.3) 84 µm (15×6 µm z-step) xyz projection. Sagittal slices corresponding to Allen Brain map slices A) 15-29 and C) 11-15 Scale bar indicated in figure.

**Figure 4.**
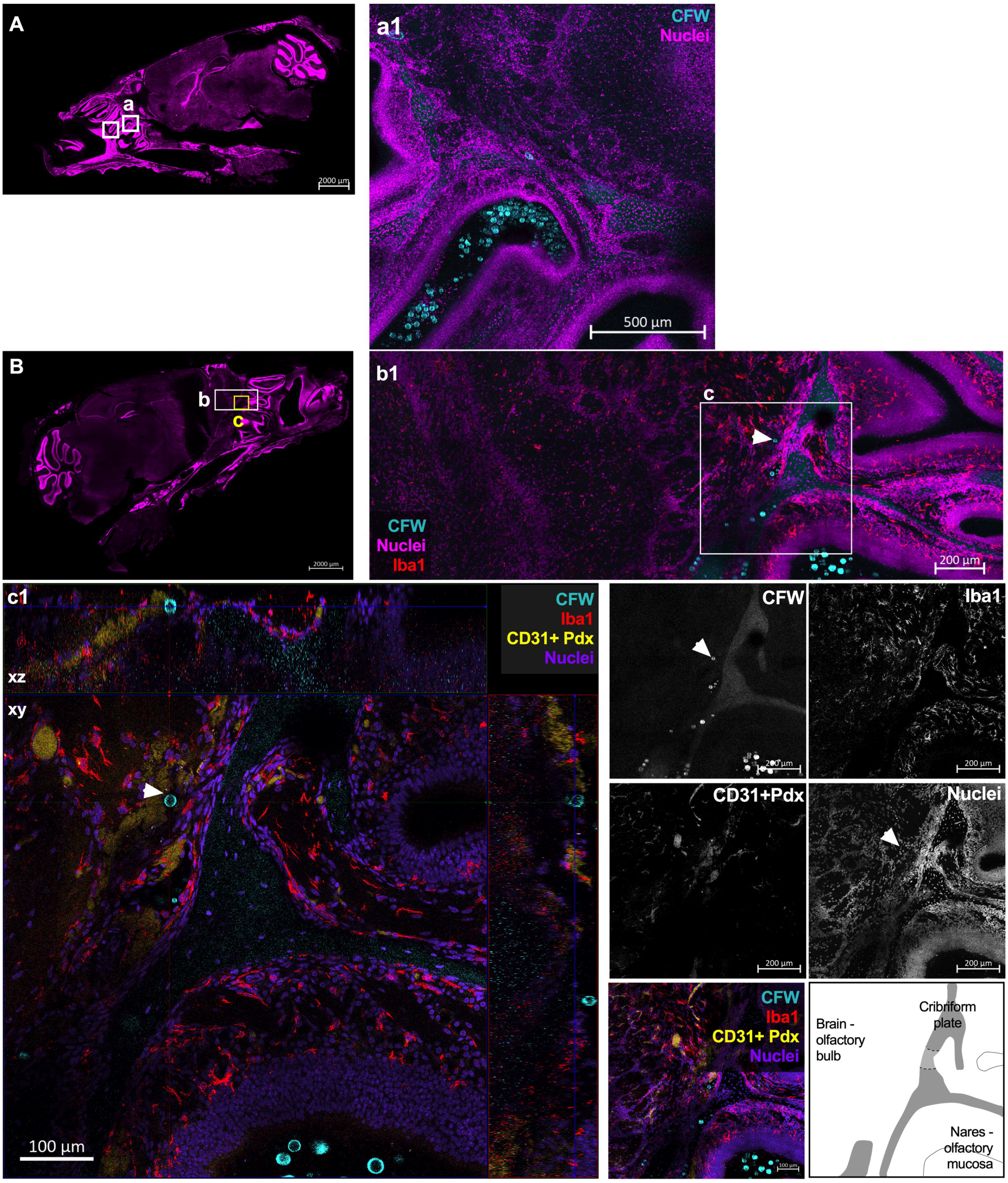
Presence of titan cells in mouse brain at 7 days post-intranasal infection, located near the cribriform plate in mouse brain as well as persistence of fungi in mouse nares. C. *neoformans* fungi, including titan cells, are present in the olfactory bulb side of the cribiforrn plate 7 dpi and continue to be abundant in upper airways and attached to ollfactory mucosa. **A-B)** C. *neoformans* fungi found in nares and ethmoturbinates(boxes). **a1)** abundant cryptococci, including titan cells in nares. **b1-c1)** C. *neoformans* cells in brain above cribriforrn plate, as well as yeasts in nares just below cribiforrn plate. **c1)** Titan cell located in brain olfactory bulb (white arrow, 17.79 µm diameter), above the cribriforrn plate. Images shown are **A-B)** single plane of skull sections, **a1)** maximum intensity projection comprising 68 µm (4 µm x 18 z-steps) deep, **b1-c1)** xyz projection with depth of 145.51 µm (2.35 µm x 63 z-steps) and maximum intensity projections of same area with a depth of 28.16 µm (2.35 µm x 13 z-steps). Scale bar indicated in images. For A-C, fungi cell wall CFW in cyan, nuclei in magenta, Iba1 in red, CD31+Pdx in yellow, with single colors images in greyscale.

### *C. neoformans* present in mouse brain parenchyma 7 days after intranasal infection

Our previous work showed that viable cryptococci could be found in mouse brains as soon as 3-24 h after intranasal infection^6^, and persist for the duration of infection^6,18^. To characterize brain invasion dynamics, clarified skulls were co-immunolabeled for blood vessels using two abundant endothelial markers, CD31 and podocalyxin (CD31+Pdx) ^19^. These sections were also immunolabeled for Iba1, a marker of microglia (brain resident macrophages)^20,21^. We confirmed specificity of these staining via single color controls (SFig.4). We also confirmed that in these sections Iba1 staining co-localized with GFP expression in CX3Cr1^GFP/+^ mouse brains, as CX3Cr1^GFP/+^ mice are frequently used to label and identify microglia in imaging and flow studies (SFig.5). Both Iba1 and GFP staining showed the characteristic microglia morphology (SFig.4 and SFig.5). Imaging and inspection of brain parenchyma from mice culled 7 days after intranasal infection showed one instance of cryptococci in the brain at 7 dpi (Fig.4B), located in the olfactory bulb above the cribriform plate. This cryptococci cell was not associated with microglia, as shown by staining of microglia Iba1^+^ cells (Fig.4B-panel c.1). Consistent with our previous observations^6^, cryptococci has already dissemination to murine brain at 7 dpi, albeit at low frequency.

### **Cryptococcus neoformans** present in the bloodstream at 3 and 7 days after intranasal infection

Several studies have shown viable cryptococci in the blood of infected mice as early as 24 hpi and up to 7 dpi in spleen and lymph nodes, which was interpreted to suggest that cryptococci disseminate from the lung to the brain via the lymph nodes, carried by antigen-presenting cells^22^. However, other works showed that cryptococci can adhere to human lung epithelial-derived A549 cells^9,23^ and other airway immortalized cells^24^. Given the capacity of cryptococci to adhere and traverse epithelial and endothelial layers, we reasoned that an additional route to the bloodstream may exist: yeasts in lung alveoli, may traverse the alveolar epithelial and capillary walls, and from here be carried in the bloodstream to other tissues, including the brain. Based on this hypothesis, we attempted to visualize yeasts directly in the vasculature of the lungs. Lungs were harvested 7 d after intranasal infection (Fig.5), stained for fungi, and this time counterstained with CD31+Pdx to label blood vessels, and with EpCAM to label airway epithelium. As expected, imaging of lungs revealed abundant cryptococci distributed through alveoli (Fig.5A) and other larger airways (Fig.5a2). We also detected 3 instances of cryptococci located in major blood vessels, identified via CD31+Pdx staining (Fig.5a3), presumably after traversal of alveoli into blood vessels. We note that we did not observe host nuclei adjacent to yeast cells, which suggest yeasts in the blood stream are free yeast, and not located within phagocytes as posited by the Trojan-horse mechanism^22,25^. To confirm viable cryptococci can be found in murine bloodstream in the first few days after intranasal infection, we additionally quantified CFU in blood extracted via cardiac bleeds (Fig.5B). We confirmed cryptococci can be found in the bloodstream as early as 3d post intranasal infection, in agreement with previous work^22^. Taken together, this suggests that cryptococci can access the bloodstream via direct traversal of airway epithelial layers in the lungs by free yeasts, in addition to being indirectly trafficked via lymph nodes by a Trojan-horse mechanism.

**Fig 5.**
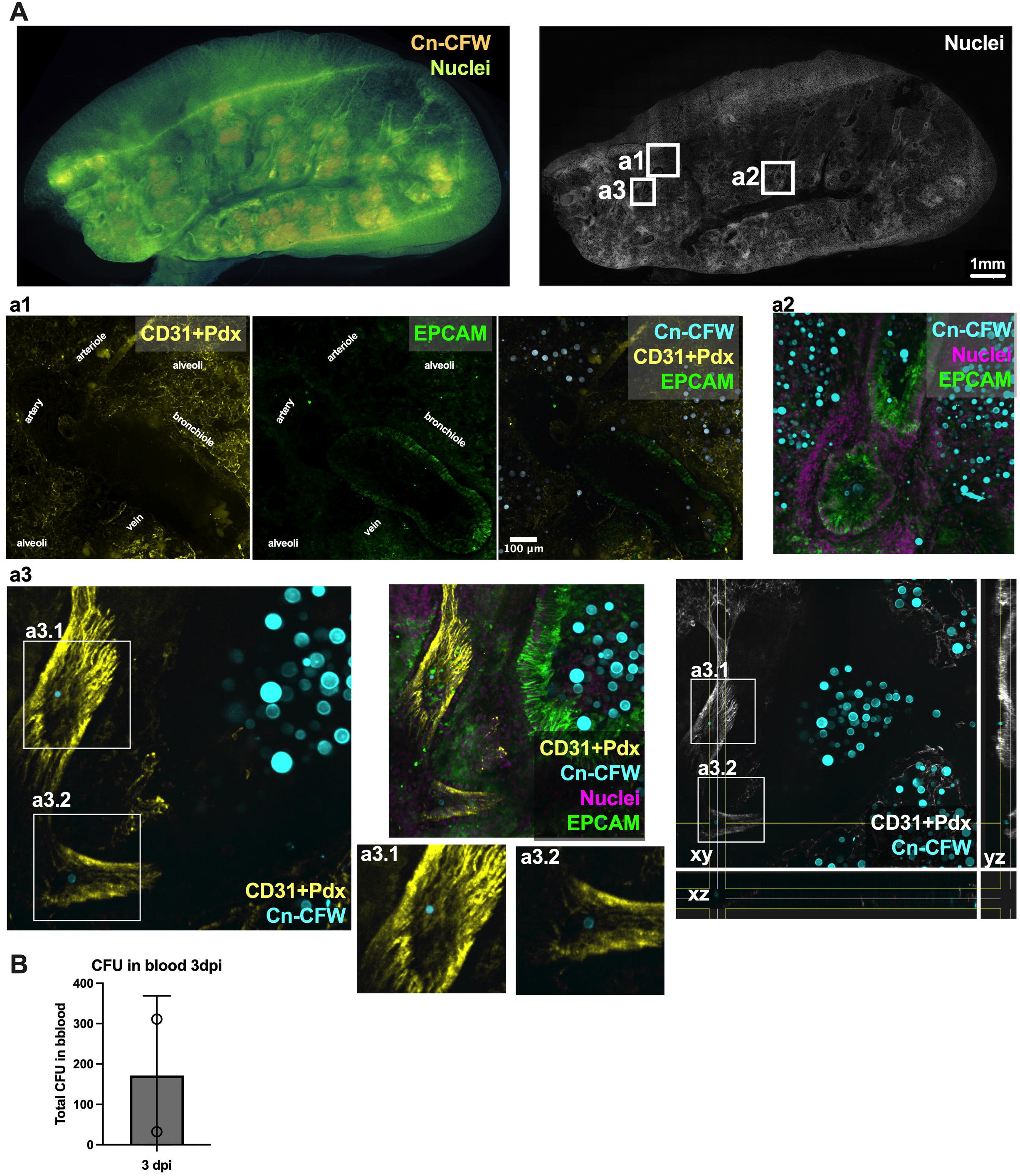
Cryptococci from alveoli access bloodstream in lungs at 3 and 7dpi. **A**) Whole lung section map allows visualization of fungi in lungs (left) followed by confocal imaging (right) showing cryptococci situated in alveoli (A-a1). A significant portion of cryptococci found in EpCAM+ airways (a2-a3), particularly in bronchioles and terminal bronchioli, and rare instances of cryptococci found in blood vessels (a3), identified by CD31+Pdx endothelium staining. B) Viable fungi can be found in bloodstream (cardiac bleed) after intranasal infection at day 3. Images are maximum projections from A) 18×25 µm z-step) a1) 23×3 µm z-step, a2) 14×13 µm z-step, a3)12×13 µm z-step, and orthogonal view in right panel. Data from from CX3Cr1^GFP/+^ (A) and C57bl/6J (B) mice intranasally infected with 5×10^5^ CFU of mCardinal H99. Data represent individual mice. Scale bar indicated in images. For A, fungi cell wall CFW in orange, nuclei in green-blue, channels merged via transparency overlay. For a1-a3, fungi cell wall CFW in cyan, nuclei in magenta, CD31+Pdx in yellow (white in xyz projection), EPCAM in green, with single colors in grayscale.

### *Cryptococcus neoformans* is distributed through the brain, has crossed BBB, and associates with Iba1 microglia within the 24 h post systemic infection

To understand the events associated with *C. neoformans* invasion of the brain, we switched to an intravenous infection route that bypasses the airway stage and initiates more rapid – and therefore experimentally tractable– brain invasion in mice. Our finding of cryptococci in blood supports intravenous injection is a valid experimental model. To enable comparison of different *C. neoformans* strains, we infected CX3Cr1^GFP/+^ mice (C57Bl/6J background) and also C57bl/6J mice with fungal strain H99E or mCardinal-KN99α (mCardinal, data pooled in Fig.6-7, SFig.6). At 24 hpi, both *C. neoformans* strains were abundant and diffusely distributed throughout the brain (mapped in Fig.6A-B), consistent with dissemination through the bloodstream^22^. We found >74% (CI:65%-84%) of cryptococci occurred in clusters (>2yeasts) instead of singlet or doublet cells (Fig.6B). At 24 hpi in brain, cryptococci ranged from ∼4-6 μm in diameter, and we did not detect titan cells nor fungal cells smaller than 3 μm in diameter in the brain parenchyma (see representative examples in Fig.1C-F, and in Fig.7), in contrast to rapid induction of titan cells in upper airways, as shown previously).

**Figure 6.**
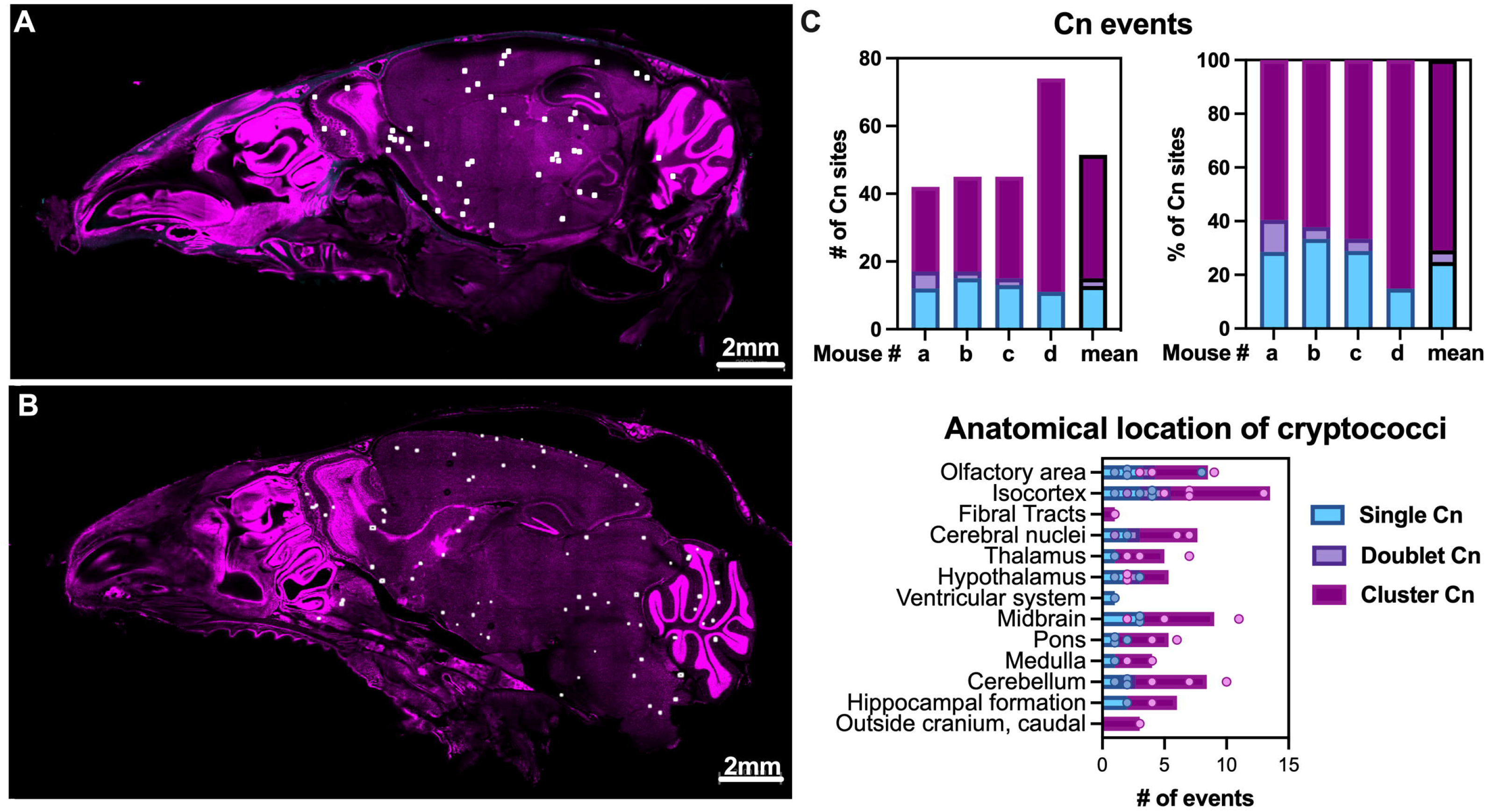
C. *neoformans* locations after intravenous infection show a dispersed pattern, consistent with bloodstream dissemination and passive arrest in cappilaries. *C. neoformans* locations dispersed through the skull, most frequently found as clusters of >2 fungi, indicating either multiple cells traversing at same location or that fungal cells are already replicating in tissue. **A-8)** Two skulls sagital section of C57bl/6J mice, with white dots indicating locations of *C.neoformans.* C) Quantification of single, doublet and clusters (>2 fungi, representative images in Fig. 1). Graphs quantify 4 mice: two males C57bl/6 each with 225 µm-thick sagital section (26×9 µm z-steps) and two female CX3Cr1GFP/+ 140-150 µm-thick (5×35 µm z-steps, 6×30 µm z-steps), 1 day after tail vein i.v. with 5×10^5^ CFU of H99E and mCardinal strain, respectively. Images shown from Images from two males C57bl/6, sagital cuts corrresponding to slices **A)** 8-14 and **B)** 13-19 of Allen Brain Atlas. **C)** Individual mice (mouse# in tops graphs, and individual dots in bottom graph) and and mean of all mice shown. CFW in cyan, nuclei in magenta.

**Fig. 7.**
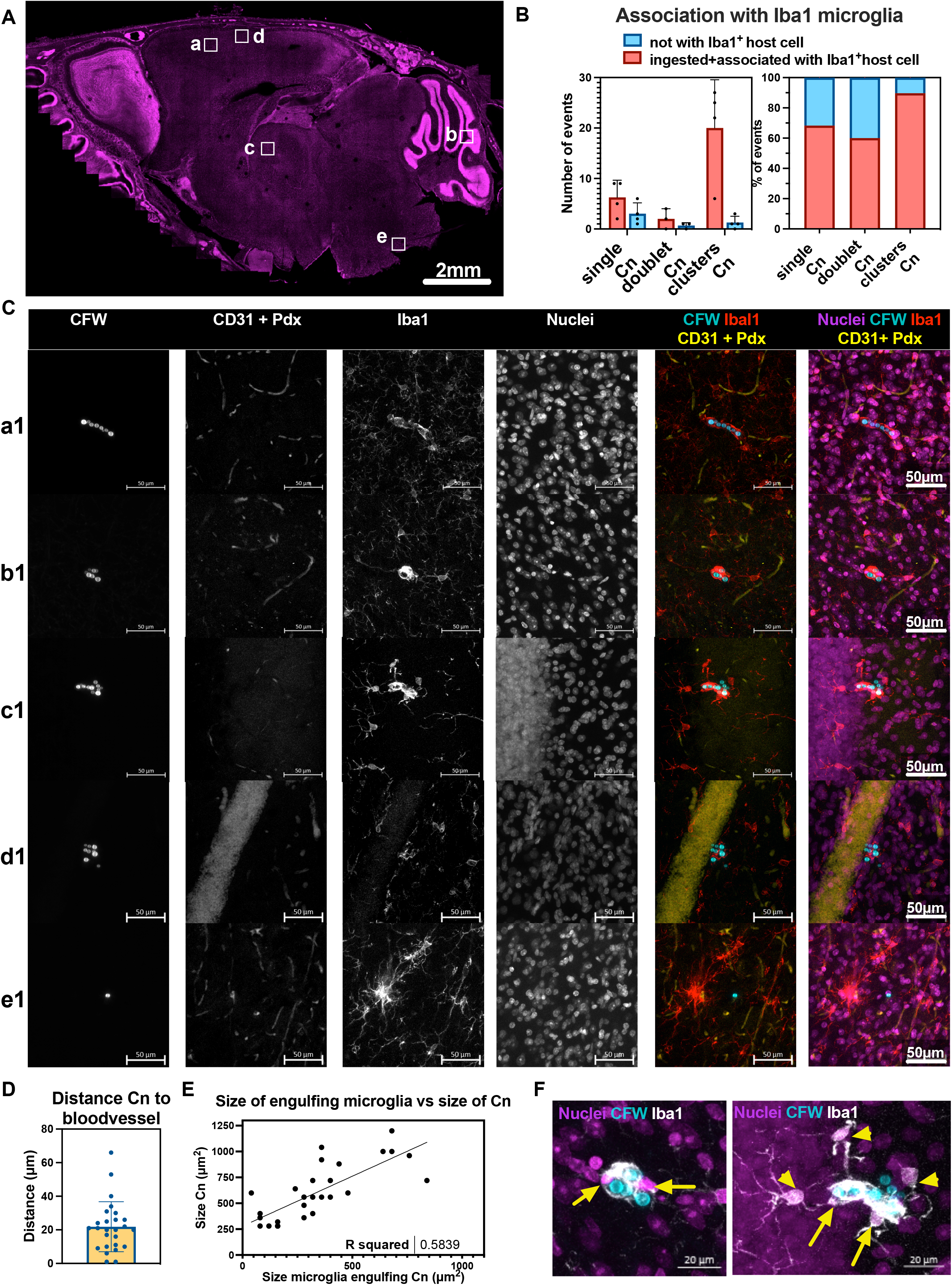
Traversal of BBB by *C. neoformans* leads to association with Iba1^+^ cells within 24hpi. Association of fungal cells with Iba1^+^ cells, the brain resident microglia, including border-associated microglia in mouse brains, as early as 24hpi. **A)** Representative image of a skull with location of *C.neoformans*. **B)** Quantification of microglia association with Cn. **C)** Representative images of cryptococci clusters associated with Iba1^+^ cells, and in proximity to CD31+Pdx^+^ blood vessels, with panel e1 showing 1 cryptococci not associated with Iba1^+^-cells. **D)** Distance of cryptococci to closest bloodvessel, stained with CD31+Pdx. **E)** Area occupied by microglia increases with size of cryptococci cluster with **F)** showing magnifications of b1 and c1 to show multiple nuclei of microglia (yellow arrows) and touching processes of neighboring microglia (yellow arrowheads). For B) data from n=4 mice, datapoints correspond to individual mice, 1 skull section analyzed per mouse, 24hpi i.v. infection with 5×10^5^ of strain mCardinal and H99E in two CX3Cr1^GFP/+^ female mice and two C57bl/6J male, respectively. For D-E) datapoints represent each cryptococci cluster, ROI randomly selected from 2 mice sections, n=26 total. Images are maximum projections of A) 140 µm (5×35 µm z-steps), corresponding to Allen Brain map sagital sections 13-18; a1) 75 µm (76×1 µm z-steps; b1) 45 µm (46×1 µm z-steps); c1) 81 µm (82×1 µm z-steps); d1) 74 µm (75×1 µm z-steps); e1) 69 µm (70×1 µm z-steps). Scale bars in images. [see SFig. 5 for representative images from C57bl/6J mice, and SFig. 6-7 for xyz projections of b1 and c1]. For A-C, fungi cell wall CFW in cyan, nuclei in magenta, Iba1 in red, CD31+Pdx in yellow, with single colors in greyscale. For F, CFW in cyan, nuclei in magenta, Iba1 in greyscale/white.

To decipher whether *C. neoformans* was present in brain blood vessels only or had crossed the blood vessels into the parenchyma at 24 hpi, we co-immunolabelled brain sections with Iba1 to label microglia and CD31+Pdx to label the vascular endothelium. We mapped fungi in brain tissue, followed by higher-resolution imaging to quantify association with Iba1-microglia and with blood vessels (Fig.7). We observed the majority of cryptococci (80%) were fully or partially encased by Iba1^+^ cells (Fig.7A-C, with representative examples in panel a1-e1 and SFig.6-8 and SVideos). Most cryptococci were adjacent to CD31+Pdx vessels (Fig.7C), with a mean distance to the closest blood vessel of 21.8 μm (CI: 15.8-27.9 μm). Only 20% of fungi were not associated with Iba1^+^ cells (example in panel e1 in Fig.7C), and these may reside in the brain parenchyma associated with host cells such as astrocytes as reported by others ^26,27^ (not labelled in our experiments) or as freely proliferating yeast in the perivascular and parenchymal space, as observed before^28–30^. We did not detect recruitment of peripheral phagocytes into the brain since GFP^+^,Iba1^-^ cells were not observed in the brain parenchyma of CX3Cr1^GFP/+^ mice, which is consistent with >14d delay in recruitment of peripheral immune cells to the brain in a systemic model of infection^31^.

Observation of microglia morphology showed microglia cells were larger following engulfment of larger cryptococci (Fig.7E) and we noted the presence of multiple host nuclei in the Iba1^+^ cluster surrounding *C. neoformans* (Fig.7F). Inspection of the morphology of fungi-associated Iba1^+^ cells indicated these microglia assumed an amoeboid-like morphology, with fewer ramified processes ^32^ (Fig.7C and F). Instances of neighbouring microglia (Fig.7F) extending processes towards cryptococci-containing microglia were noted, suggesting communication from infected microglia to neighbouring microglia ^32^. The observed enlargement of Iba1^+^-microglia may occur by migration and fusion of neighbouring microglia upon infectious stimuli; alternatively microglia in response to inflammatory stimuli *in vitro* become multinucleated due to cell proliferation with failed cytokinesis ^33^. We did not observe nuclear morphology suggestive of active proliferation by microglia, but our previous work showed that *in vitro* phagocytes and *in vivo* alveolar macrophages proliferate in response to cryptococcal infection^34,35^. These aspects will be dissected in future studies.

Overall, our data shows that within 24 h of systemic murine infection, *C. neoformans* crosses BBB within the first hours after arrest in brain capillaries. Since the majority of cryptococci are in clusters, we propose 2 possible explanations: traversed cryptococci start to proliferate in the first 24 hpi, and possibly very soon after traversal of the BBB; alternatively, it is possible that cryptococci traversal creates a transient breach in the BBB, which can be exploited by subsequent cryptococci arrested at the same capillary site. Soon after traversal, cryptococci in parenchyma are phagocytosed by brain resident microglia, prior to recruitment of peripheral monocytes (summarised in Fig.8). Generally, microglia enlarge and may show multiple nuclei when interacting with multiple cryptococci. In summary, our data make an important first contribution towards characterising the myriad cellular interactions required for *C. neoformans* invasion of the mammalian brain.

**Fig. 8.**
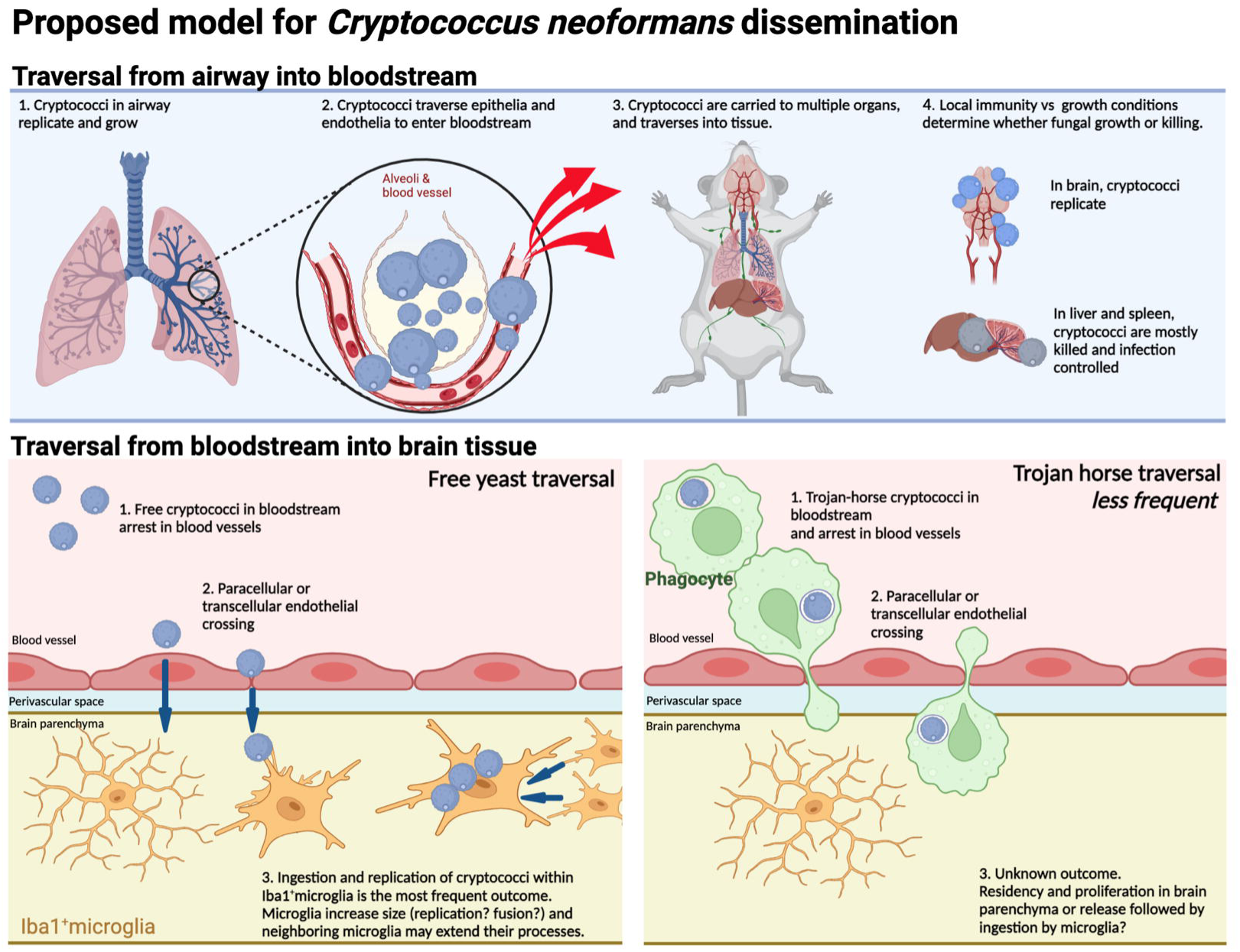
*Cryptococcus neoformans* traversal of barriers culminating in brain parenchyma invasion. Figure created in Biorender.

## Discussion

In this work, we harnessed clarified tissue sections to image host-pathogen interactions in tissues at subcellular resolution. Thick sections (200 μm imaged) provided a step-change in analysing cryptococcal-tissue interactions and is easily adaptable to a wide range of microscopes. Thus, the imaging and analysis pipelines we established may be very easily adapted in other laboratories. We confirmed that crossing the BBB is performed by *C. neoformans*, and show for the first time in mice that microglia, the brain resident macrophages, rapidly ingest traversed fungi, all within 24 hpi after systemic infection. This approach also provided new information on interactions of *C. neoformans* with murine hosts, including observation and quantification of titan cells *in situ*.

Whilst we and others had cryptococci in upper airways of mice^36,37^, we report for the first time that titan cells are abundant in airway turbinates of mice. Here, titan cells were found at high frequency in the first 24 hpi and at least up to 7 dpi, which indicates very strong titan-inducing conditions in airway turbinates. To our knowledge this is the fastest rate of titan cell formation, as titan cells are rare in the lung at 24hpi but reach ∼20% of fungi by 3 dpi^38^. Additionally the presence of *Cryptococcus* in upper respiratory tract is common in several animals, particularly dogs and cats^39^, koalas^40,41^ and ferrets^42^, and indicates either asymptomatic carriage or a symptomatic upper respiratory tract infection, which can progress to invasive infection^40^. Thus, it is probable titan cells occur in airways of most animals, and it is important to investigate their contribution to veterinary infection and disease. Our observation of significant fungal burden in airways, together with an event of mucosal invasion in upper airways, suggests upper airway residency may be a relevant site of *C. neoformans* infection, facilitating invasion of extra-airway sites either as a reservoir for fungal cells or as an additional site of access to the bloodstream via crossing of the nasal mucosa.

The data on nasal carriage of cryptococci in humans is sparse compared to other animals. Nevertheless, asymptomatic carriage of cryptococci is possible: one case report describes that after a pet ferret was diagnosed with cryptococcosis, its human owners showed positive cultures from nasal swabs, despite negative serum antigenemia^42^. Thus, whilst colonization of human noses in immunocompetent individuals is possible, this is still insufficient data on frequency of nasal carriage in healthy immunocompetent humans and whether this is transient or long-lasting. Additionally, cryptococci can be detected in the upper respiratory tract of patients with symptomatic cryptococcal disease. One study in Lisbon, Portugal observed abundant fungal cells in olfactory mucosa via histopathological analysis of the autopsies of patients who succumbed to AIDS-associated cryptococcosis^43^, whilst a study in Nonthaburi, Thailand recovered viable cryptococci from the nasopharynx of individuals recently diagnosed with AIDS-associated cryptococcal meningitis. Whilst the presence of cryptococci in the nose may be attributable to the high fungal burden in these patients, it remains to be determined whether presence of cryptococci in upper airway of humans occurs during early disease of humans, and whether, if at all, contributes to pathogenesis and/or persistence in humans.

Bloodstream dissemination of cryptococci is widely accepted, as it is consistent with a diffuse and broad distribution of *C. neoformans* throughout the brain, in close proximity to blood vessels, observed in in humans post-mortem brains^44,45^ and that we and other characterized in murine models^29,30,46^. Additionally, murine studies can detect blood borne cryptococci after intranasal inoculation^22^. Previous studies showed rapid trafficking of yeast and spore-derived yeasts into murine lung-draining lymph nodes, as early as 24 hpi^22,47^, and posited that escape from lung into lymph node and then into bloodstream provided a route to the murine brain. In this study, spores had quicker dissemination kinetics than yeast particles, ^22^, via mechanisms yet unknow. Here, we observed that large blood vessels in lungs contain free cryptococci, presumably after crossing alveolar airway and endothelial layers. Thus, our data is indicative, but not conclusive, of free yeast traversal being the predominant mechanism, and that direct traversal of lung alveoli may be an alternative and significant route towards bloodstream dissemination. Our hypothesis is supported by work showing cryptococci can adhere to human lung epithelial A549 immortalized cells^9,23^ and other airway immortalized cell lines^24^. An open question is how *C. neoformans* traverses these tissue layers to reach the bloodstream. Strategies to escape from upper and lower airway, contribution of different fungal particles and morphotypes^22,48^, as well as the relative contribution of lymphatic dissemination via direct angioinvasion into the circulating bloodstream, remain to be determined.

Rapid traversal of BBB by *C. neoformans* was previously detected via intravital imaging to show that tail vein injection of particles of a certain size, such as fungi and inert polystyrene beads, leads to passive trapping of particles in small brain capillaries. Six hours after injection live fungi, but not killed fungi nor beads, traversed capillaries into the brain parenchyma^7,8,49^, indicating an active process of crossing by the pathogenic fungus. We also observed rapid crossing of BBB by *C. neoformans* followed by close interactions with Iba1^+^ microglia, including ingestion of cryptococci, and presence of fungal clusters suggesting replication and growth within microglia or in the brain parenchyma. Our observations are in line with previous work determining that the brain niche is favourable to cryptococci growth, due to a combination of localized immune features and to favourable nutrition, for example an abundance of manitol^12^. We extend these results by showing early ingestion of fungi by Iba1^+^ cells in the very first day of infection, after endothelial crossing into brain parenchyma. Rapid association of cryptococci with brain-resident macrophages is in line with recent publications. First, ingestion of a small percentage of cryptococci by microglia 4 d post infection in the developing brains of zebrafish larvae^50,51^. Second, flow cytometry analysis of immune cells in mouse brains, 7 d after intravenous infection, showed cryptococcal association with phagocytes in mouse brains, even after tissue dissociation, albeit a significant percentage of fungi are extracellular ^28^; at this stage, fungi-containing cells were either microglia or phagocytes infiltrating from periphery. Third, previous studies show perivascular yeasts in BBB, either in free form or associated with phagocytic cells, 3 to 7 d after retro-orbital or tail vein inoculation^30^. In contrast, two other studies find a significant percentage of fungi in mouse brains are extracellular. Up to 18h after intravenous infection, the majority of yeast in brain lysates are extracellular, with less than 10% associated with leukocytes^10,52^. We posit that this discrepancy arises due to technical constraints: protocols which disrupt tissue may disrupt or discard clusters formed by microglia and fungi. This would suggest that our data is more reflective of true tissue interactions. Whilst the percentage of phagocyte-associated vs free fungi may vary during the course of infection, soon after BBB crossing the majority of cryptococci are interacting and in close proximity to microglia and trigger localized responses in adjacent microglia cells. Still, our studies pave the way to study localised, spatially resolved host-fungal interactions underpinning invasion, such as determining the relative contribution of Trojan-horse traversal ^25,53,9,52^, the contribution of Mpr1^11^ and interaction between hyaluronic acid in capsule and CD44 in endothelial cells to brain tissue invasion^54^.

We observed apparent association between multiple amoeboid microglia to ingest clustered cryptococci. Expression of an amoeboid morphology is commonly associated with the inflammatory activation of microglia in several pathological processes^55^. This immune activation will likely occur at all stages of infection, as was previously observed in a model of late cryptococcal meningitis, following intracerebral infection of mice^46^. Amoeboid microglia were also observed after *Streptococcus pneumoniae* infection^56^; in contrast, ramified microglia are still observed in the first hours after *Toxoplasma gondii* infection^57^, demonstrating an interplay between neuro-immune response and invading microbe. At this stage there is no detectable recruitment to infection sites of circulating monocytes. Further characterization is needed to drill down into the functionality of these Iba1^+^ cells. Whilst Iba1 is a well-accepted microglia marker, immune populations are now recognized as more complex and heterogeneous even within the same organ. Recently, transcriptomic and developmental profiles showed Iba1^+^ cells in brain are “true” microglia, parenchyma-resident macrophages which can migrate to vessels in response to invading stimuli, but a second population of Iba1+ cells are brain border associated macrophages, and these subsets have subtle but important functional distinctions^55,58^.

One noteworthy observation is that sites with permeable capillary beds, such as the liver and the spleen, show reduced burden of *C. neoformans*, and yet the brain, seemingly well-protected by the multi-layered BBB, is effectively colonized. *C. neoformans* trapped in liver sinusoids after systemic infection was ingested by Kupfer cells, the liver-resident macrophages, and fungal burden controlled in the first few hours post –injection^49^. The corollary of our studies is that tissue barriers, including BBB, are not fully impermeable to pathogens, and bloodstream permeability is not a major determinant of tropism over the course of infection, at least for *C. neoformans*. Instead, after seeding of fungal pathogen in several organs, cryptococcal tissue tropism is likely most determined by the underlying tissue-specific immunity and by the pathogen‘s adaptations to the specific nutritional conditions of the tissue, reminiscent of the “seed and soil” hypothesis by S. Paget^59^.

Overall, we show here a high-content, high-resolution method to study fungal-host pathogens, including fungal morphological analysis and tissue-immune interactions. This method potentiated several observations: we confirm the presence of abundant Titan cells in airway turbinates of mice, as we reported before^6^, we observed for the first time *C. neoformans* in the lamina propria of murine turbinates, and confirm presence of cryptococci in bloodstream of mice. Further we show that in the early stages of brain invasion, similar to what occurs in lungs, *C. neoformans* associates rapidly with tissue-resident phagocytes, in this case Iba1^+^ cells. Our work unveils early events in *C. neoformans* invasion of mammals and new insights into cryptococcal disease.

## Methods

### Fungal strains

We used *C. neoformans* H99E, originating from JE Lodge laboratory and deposited into Fungal Genetics Stock Centre, for the majority of experiments. Strains *ste50*Δ and *cac1*Δ were obtained from deletion library, created by the Madhani laboratory^60^, through Fungal Genetics Stock Center. Strain H99-mCardinal (CnLT0004) was a gift from Edward Wallace and Laura Tuck; mCardinal is derived from KN99α to expresse the mCardinal red fluorescent protein^61^, codon-optimised for *Cryptococcus*, and integrated into genomic safe haven 4 ^62,63^ with RPL10/CNAG_03739 promoter and terminator, and a NAT resistance cassette. Cryptococci were grown from frozen 10% glycerol stocks on yeast extract-peptone-dextrose (YPD) agar plates for 2 days at room temperature, followed by culturing overnight at 37 °C, 180 rpm, in YPD broth. Cryptococcal cell suspensions were counted in hemocytometer and diluted to the appropriate density.

### Mouse infections

C57BL/6J male mice, aged 8 to 12 weeks, were purchased from Charles River Laboratories, UK and infected with 5 x 10^5^ CFU, unless otherwise specified, in sterile phosphate buffered saline (PBS, Oxoid, BR0014G). Intranasal infections were performed by placing 25 µl of yeast suspension into the mouse nares under isoflurane anaesthesia. Intravenous infection was performed with 5 x 10^5^ CFU in 100 µl via tail vein injection. Mice were monitored every 6 h for the first 24 h, and then daily, for deterioration in health. We also imaged noninfected (sentinel) mice for tissue morphology, immunolabel, and dye staining controls, including for CFW and antibody staining specificity. CX3CR1^GFP/+^ mice were obtained from University of Exeter colony, a kind gift from Jon Witton and Peter C Cook. All experiments were approved by the University of Exeter under protocol P79B6F297, and P6A6F95B5.

### Tissue extraction and fixation

Mice were culled via cervical dislocation. Skin was removed, skull and thorax was opened to remove lungs. For samples intended for skull imaging, the skin, lower jaw, tongue, and attached skull muscles, were removed. When needed, cardiac bleeds were performed under isoflurane anaesthesia. All tissues were fixed for 48 h in approximately 20-fold volume of 4% formaldehyde at room temperature with gentle agitation in a rotating shaker. Tissues were then rinsed several times in PBS containing 0.02% azide and stored at 4℃ until further processing. Unless otherwise noted, 0.02% azide was added to all PBS-based solutions to prevent microbial growth.

### Decolourisation and decalcification

After fixation, skulls were placed in decolourisation solution made with 30% dilution of CUBIC reagent 1, as in ^14^, in 0.1M PBS (CUBIC reagent 1 was prepared with 25 wt% urea (Thermo Scientific, U/0500/65), 25 wt% N,N,Ń,Ń-tetrakis (2-hydroxypropyl)ethylenediamine (Thermo Scientific, L16280.AE) and 15 wt% Triton X-100). Skulls were incubated at 37 °C for 48 h in 5 ml decolourisation solution with the solution being refreshed at least four times until it remained clear. Samples were washed twice in 5 ml PBS and placed in 40 ml decalcification solution (0.2M ETDA in 0.1M PBS adjusted to pH 8-9 with sodium hydroxide) for 72 h at 37 °C. Skulls were washed twice in 5ml PBS and kept at 4°C in PBS until further processing.

### Slicing and tissue clearing

Whole organs were submerged in up to 5 ml X-CLARITY hydrogel monomer with initiator solution. Oxygen was removed from the solution via degassing with nitrogen flow prior to organ submersion and again after submersion. Tissues were incubated at 4 °C overnight followed by 3 h at 37 °C, with gentle agitation. Tissues were washed in 5 ml PBS to remove hydrogel. Organ sections were obtained by 300 or 400 µm sagittal cuts with a vibratome. In some cases, tissues were cut before embedding in hydrogel, but we found tissue to become more stable if hydrogel-embedded was performed before cutting. All tissues were cleared with X-CLARITY following manufacturer instructions. Tissues were incubated in 2 ml X-CLARITY tissue-clearing solution at 37 °C overnight. Tissues were then cleared in an electrophoretic tissue clearing chamber (ECT, LogosBio instruments) with a current of 1.5A, circulation speed of 30 rpm, at 37 °C for 3 h, and inverted halfway through incubation. Samples were washed twice in 5ml PBS and stored at 4 °C in PBS, until further analysis.

### Staining

Organ sections were stained with CFW for 48 h with gentle rotation at room temperature prior to blocking and staining with antibodies and dyes. Tissues were blocked overnight in 1ml Fc block solution, containing 5 µg/ml 3.G2 Fc-block (BD Bioscience, 553142), 3% BSA, 0.02% azide, 0.1% Triton X-100 in PBS. Tissues were stained with antibodies and dyes listed in Table 1, in Fc blocking solution, for 48h with gentle agitation. After staining, samples were rinsed in 1 ml PBS overnight at room temperature with gentle agitation. Prior to imaging, tissues were mounted in refractive index match solution (80% glycerol in water).

**Table 1.** List of Antibodies and dyes used.

| Antigen-Fluorophore | Clone | Catalog # | Company | Staining Concentration (Dilution from supplier stock If conc not available) | Target |
| --- | --- | --- | --- | --- | --- |
| CD31-PE | MEC13.3 | 102507 | Biolegend | 5-7 µg/ml | Endothelium |
| CD31-AF647 | MEC13.3 | 102515 | Biolegend | 5-15 µg/ml | Endothelium |
| Podocalyxin-PE | 10B9 | 107930 | Biolegend | 5-7 µg/ml | Endothelium |
| AF647 – Podocalyxin | 192703 | FAB1556R | Bio-techne | 22.5 µg/ml | Endothelium |
| Iba1-AF647 | EPR16588 | AB225261-1001 | Abcam | 5 µg/ml | Iba1, microglia specific marker |
| EpCAM-PE | G8.8 | 15228839 | Invitrogen | 5 µg/ml | Lung Epithelial |
| Helix NP<sup>TM</sup> Blue |  | 425305 | Biolegend | 25 µM | Nuclei |
| Helix NP<sup>TM</sup> Green |  | 425303 | Biolegend | 25-200 µM | Nuclei |
| Propidium iodide |  | P4864 | Sigma | 20ug/ml | Nuclei |
| CFW (Fluorescent Brightener 28) |  | F3543 | Sigma | 50-100 µg/ml | Chitin in cell wall of fungi |

### Imaging and processing

For imaging tissue sections, we used two confocal microscopy systems. On a Zeiss LSM 880 Airyscan, we firstly generated a tissue outline by briefly imaging with a 10X objective (PApo 10X0.45 – dry), which optimized the imaging area for subsequent imaging in detailed with a 25X LD LCI PApo 25×/0.8 Imm Corr D objective, with oil immersion (RI=1.518). Combinations of 405nm, 488nm, 561nm and 633nm lasers were used, depending on antibody/dye combinations. In most cases, the visible beam path was set up to use a tri-main beam splitter (488/561/633) and the UV path used a-405, beam splitter. At minimum, images were acquired with a zoom of 0.8 and a pixel voxel of 0.83 µm x 0.83 µm x 9 µm with resolution increased in some regions of interest (ROI) to >0.531 µm x 0.531 µm x 6 µm. Multicolor images were obtained using a combination of spectral 32 channel GaAsP PMT and multi-alkali PMTs detectors. Antibody staining controls were performed on unstained and single stained samples, and informed acquisition and processing of images (See SFig.1,4). Zeiss Zen [Blue edition] 2.3 or [Black edition] were used for stitching images together for full maps, with a 15-20% overlap.

Some sections were imaged on a Nikon Ti2 microscope body (Nikon Microsystems) with a DragonFly505 40 µm pinhole spinning disk microscope (Andor, Belfast) operating Fusion software. UV fluorophores was excited with a 405 nm laser and the emission collected through a Semrock TR-DFLY-F445-046 filter. Green fluorophores were excited with a 488 nm laser and the emission collected through a Semrock TR-DFLY-F521-038 filter. Full configuration of microscope available at https://www.fpbase.org/microscope/ZR9nEdoko3bwnnU66QAJUE/. Fluorescence was collected on an Andor Sona sCMOS camera with 2×2 binning for large area maps with some ROI imaged with 1×1 binning. Overview skull maps were generated using a Nikon 4×/0.2NA PlanApo lambda lens, while higher resolution 3D acquisitions were acquired using a Nikon 20×/0.75NA PlanApo lambda lens. Images were stitched using 5% overlap on Fusion ClearView software. Z-stacks and step size are indicated in the figure’s legends.

### Automated measurement of fungal cell diameter

Automated 2D fungal cell segmentation was achieved by selection of an ROI, and z-stacks were transferred to ImageJ ^64^, the fungal cell wall CFW channel was extracted and normalised. Areas of certain images with high background autofluorescence (bone protrusion in nasal cavity) were cropped out prior to analysis. Loss of signal intensity with depth of tissue was corrected with xyz normalization through z-stacks in ImageJ. Manual tracings were performed on z-stacks (before sum projection), using Fiji straight or freehand line tool for cross section and object boundary tracing for interpolated diameters based on cell area. For Stardist, ROI were sum projected to produce a 2D image from the 3D z-stack, and 2D images analysed with StarDist ^65^, with probability/score threshold of 0.15, an overlap threshold of 0.40 and a boundary exclusion of 5 for single-cell area analysis. Cell diameters were then computed from the area of objects, using the formula for area of a circle = π x radius^2^.

### Data analysis

Images were extracted and figures assembled using a combination of Zeiss Zen Blue edition 2.3, Zen Black (Carl Zeiss Microscopy), Imaris (Oxford Instruments), and ImageJ (NIH, USA). Unless stated on figure legend, no image processing steps were performed. Distances of cryptococci to blood vessels were measured in 3D stacks, using manual tracing. Areas of microglia and fungi were measured using spatial stereology, which maps areas with minimal bias based on methods by ^66,67^. Graph Pad Prism was used for graphs and statistics, Biorender was used to create diagrams.

## Supporting information

SVideos

## Acknowledgements

We thank all those that provided helpful comments in the manuscript.

This work was supported by AMS Springboard Award SBF006\1024 (UK), Wellcome Trust Institutional Strategic Support Award (WT105618MA) to CC. We acknowledge funding from the MRC Centre for Medical Mycology at the University of Exeter (MR/N006364/2 and MR/V033417/1), the NIHR Exeter Biomedical Research Centre. The project was supported by ARUK Network Centre Grant ARUK-NC2021-SW. Additional work was undertaken by the University of Exeter Biological Services Unit. A.C. was supported by NIH grants AI052733-16, AI152078-01, and HL059842-19. JMH is funded by NS127076 and AI168539. We thank Edward Wallace and Laura Tuck, for the kind gift of mCardinal strain. Fungal strains collection was funded via NIH funding (R01AI100272) led by Hiten Madhani, UCSF. The funders had no role in study design, data collection and analysis, decision to publish, or preparation of the manuscript.

## Supplemental Results

We tested several automated and manual pipelines to measure the number and size distribution of cryptococcal cells, including ImageJ (StarDist macro, open access software^64^). To ensure these pipelines were accurately measuring titan cells (Fig 2D-J), we measured fungal sizes with manual cross-section to ascertain diameter of titan cells (the most commonly used method ^68–70^). We noted that automated algorithms calculate diameter from circumference tracing, and thus we also performed manual conference tracing (interpolated ∅– cell area). Both circumference tracing methods (manual and StarDist) resulted in an increased diameter, and thus a higher % of titan cells, than cross-section diameter measurements, and this needs to be considered when comparing our data to previous studies. Still, we found that manual circumference tracing and automated measurements were comparable in detecting number, and average diameter of cryptococci (Fig.2I).

To examine variations in titan cell formation *in situ*, and the accuracy of our fungal size pipelines, we required tissues where titan cells were formed and tissues were titan cells were rare. From the available deletion library, we obtained the *cac1*Δ strain, which fails to produce titan cells *in vitro*^69,71^, and as control, we used *ste50*Δ, which having undergone similar genetic engineering retains wild-type virulence and capacity to form titan cells^13^. At 5 days post infection (dpi) with *ste50*Δ (5×10^7^ CFU), mouse lungs contained a wide size range of cryptococcal cells, with an average fungal cell diameter of 8.4 μm. To ensure we had and few titan cells and abundant fungi in lungs, we imaged *cac1*Δ at 24 hpi, as titan cells are rare in lungs at 24 hpi^38^, whilst ensuring *cac1*Δ fungi were abundant and not yet cleared by host immune system. In lungs infected with *C. neoformans cac1*Δ (5 x 10^7^ CFU) and imaged 24 hpi, our pipelines detected no titan cells and all cryptococci had a cell body size below 6 μm, with some cells as small as 2-3 μm (SFig.3).

X-CLARITY can expand some tissues significantly, but this expansion is largely reversed on refractive index (RI) matching, the last step before imaging ^17,72^. To confirm our imaging pipeline was reliably detecting titan cells, we submitted YPD-grown cryptococci to the clarification protocol and observed no changes in size (data not shown).

At this point, we reasoned that clarification allows thick tissues to be imaged in other microscopes, allowing certain measurements to be made in a great range of microscopes. Thus, we imaged lung sections in a non-confocal microscope with deconvolution capabilities, such as widely available microscope Deltavision ELITE. A clarified infected lung was imaged up to 176 μm depth (SFig.3), with 2 μm z-steps. After deconvolving, removing out of focus and low intensity images (first eleven z-steps are used for deconvolution calculations and last z-steps had lower intensity of signal), images of usable quality, with enough resolution to observe fungal cells, could be obtained between z-steps 12 to 53, for a final stack totalling 82 μm depth, and an actual tissue penetration of ∼100 μm, which greatly comparable to confocal microscopy. A separate experiment shows nuclear staining can also be observed in thick tissues after clarification (data not shown). Whilst the level of detail obtained does not reach confocal resolution, our data indicates tissue clarification expands the range of microscopes that can be used to image thick tissue and to image structures approximately the size of cryptococci or mammalian nuclei (>1-4 micron); but not sufficient for structures smaller than those.

