## Supplementary figures and images for "*Cryptococcus neoforman*s rapidly invades the murine brain by sequential breaching of airway and endothelial tissues barriers, followed by engulfment by microglia"

### SFig1. CFW-nuclei controls.tiff

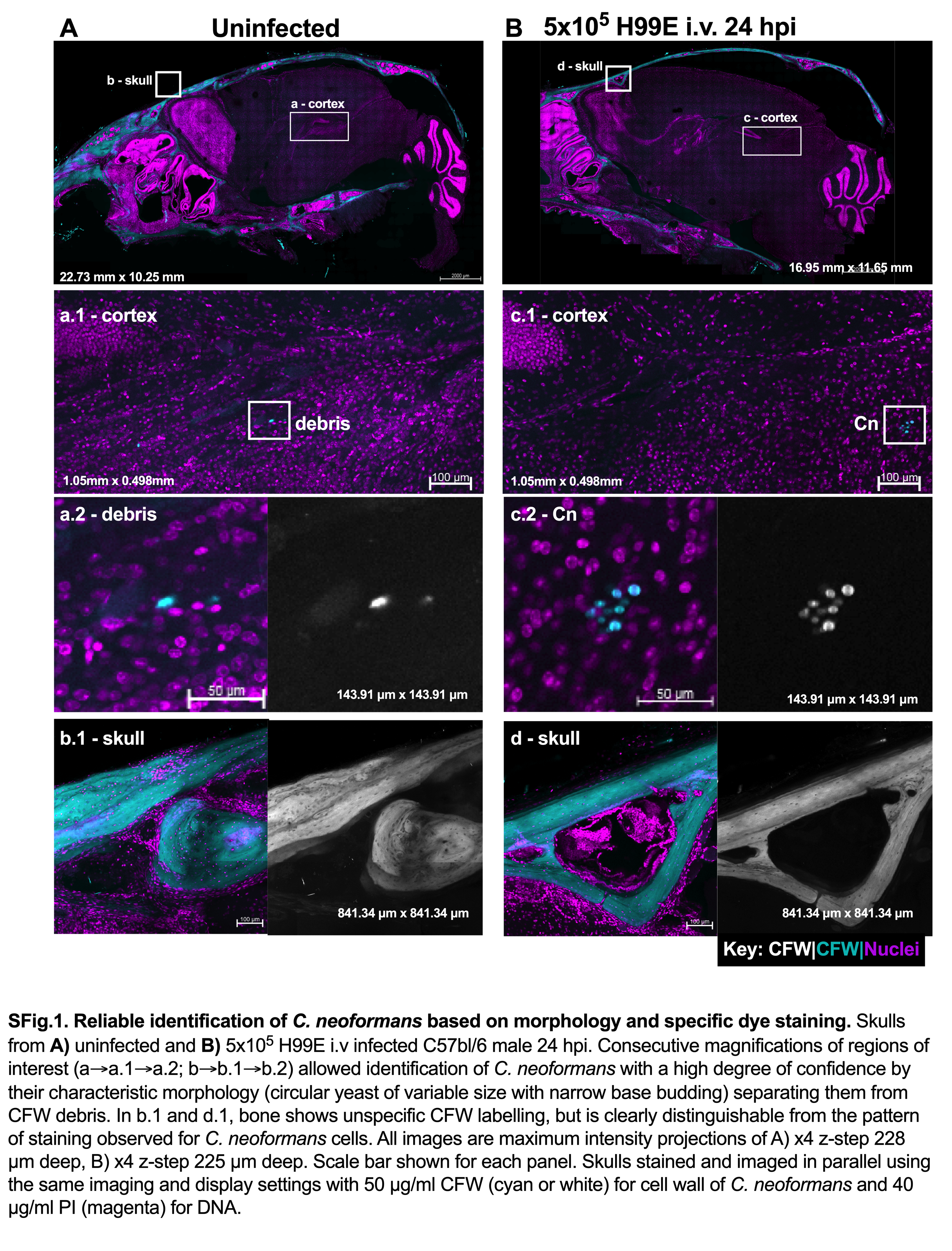

### SFig2. TitanLungVNov2023.tiff

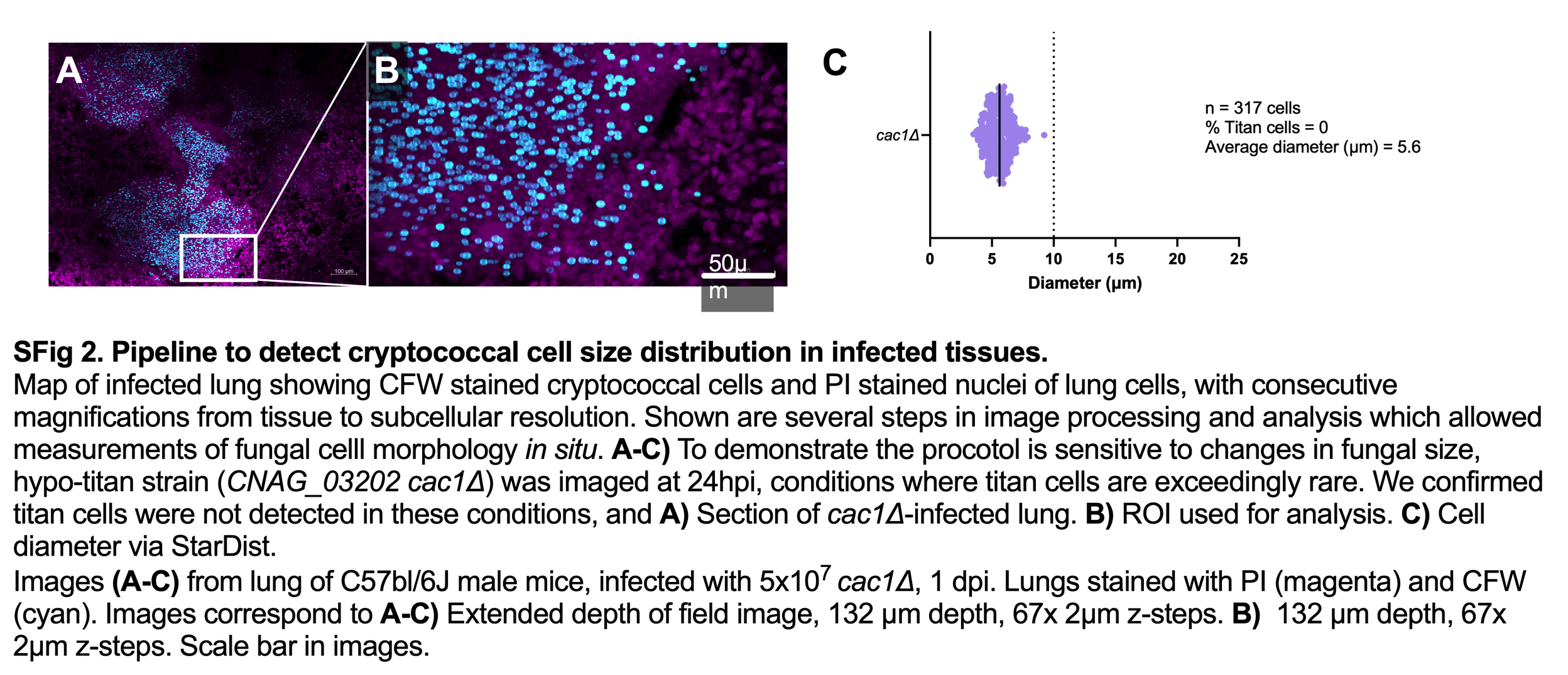

### SFig3 DV image clean.tiff

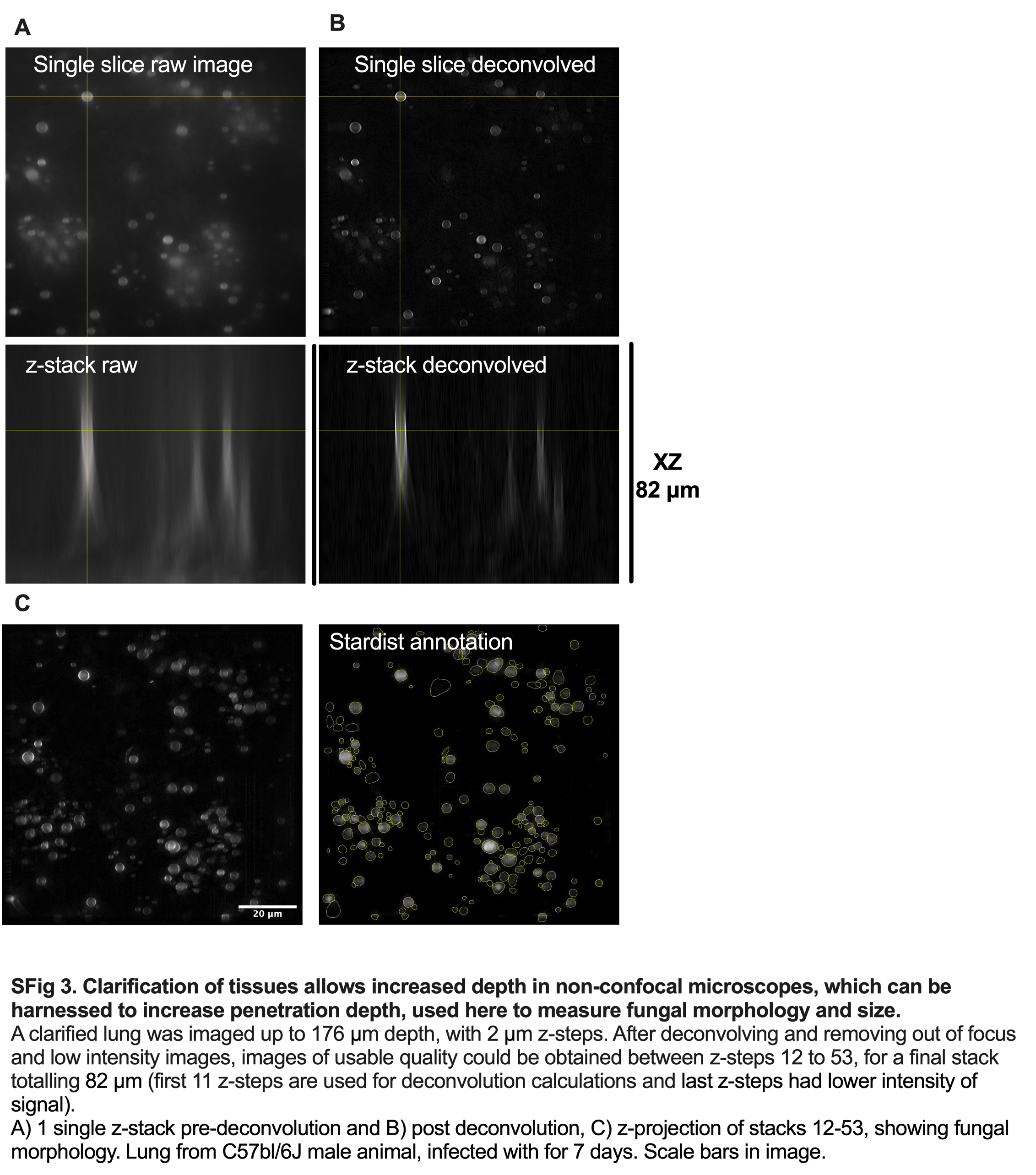

### SFig4 Single Colori.tiff

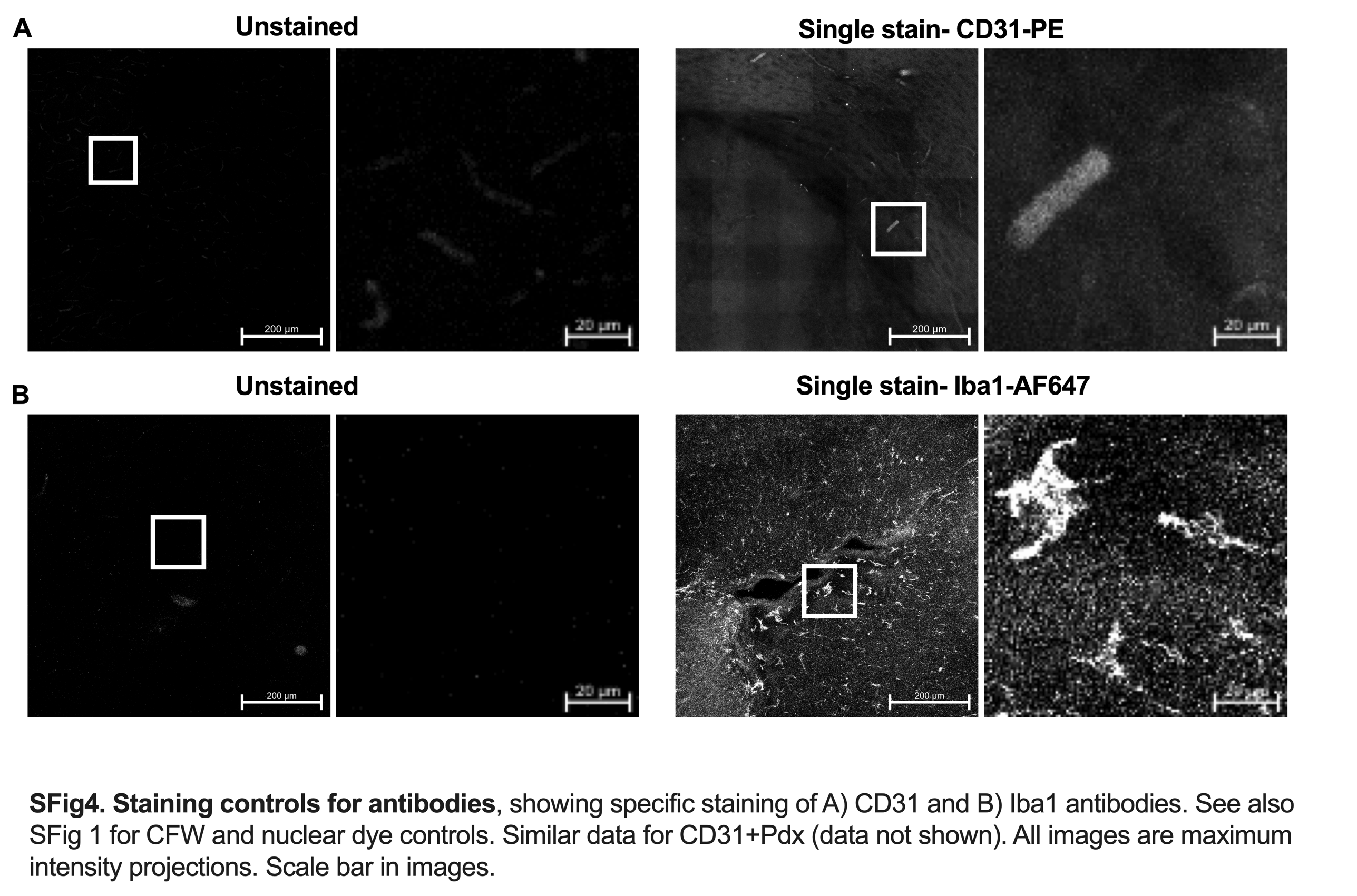

### SFig5. WT lba1.tiff

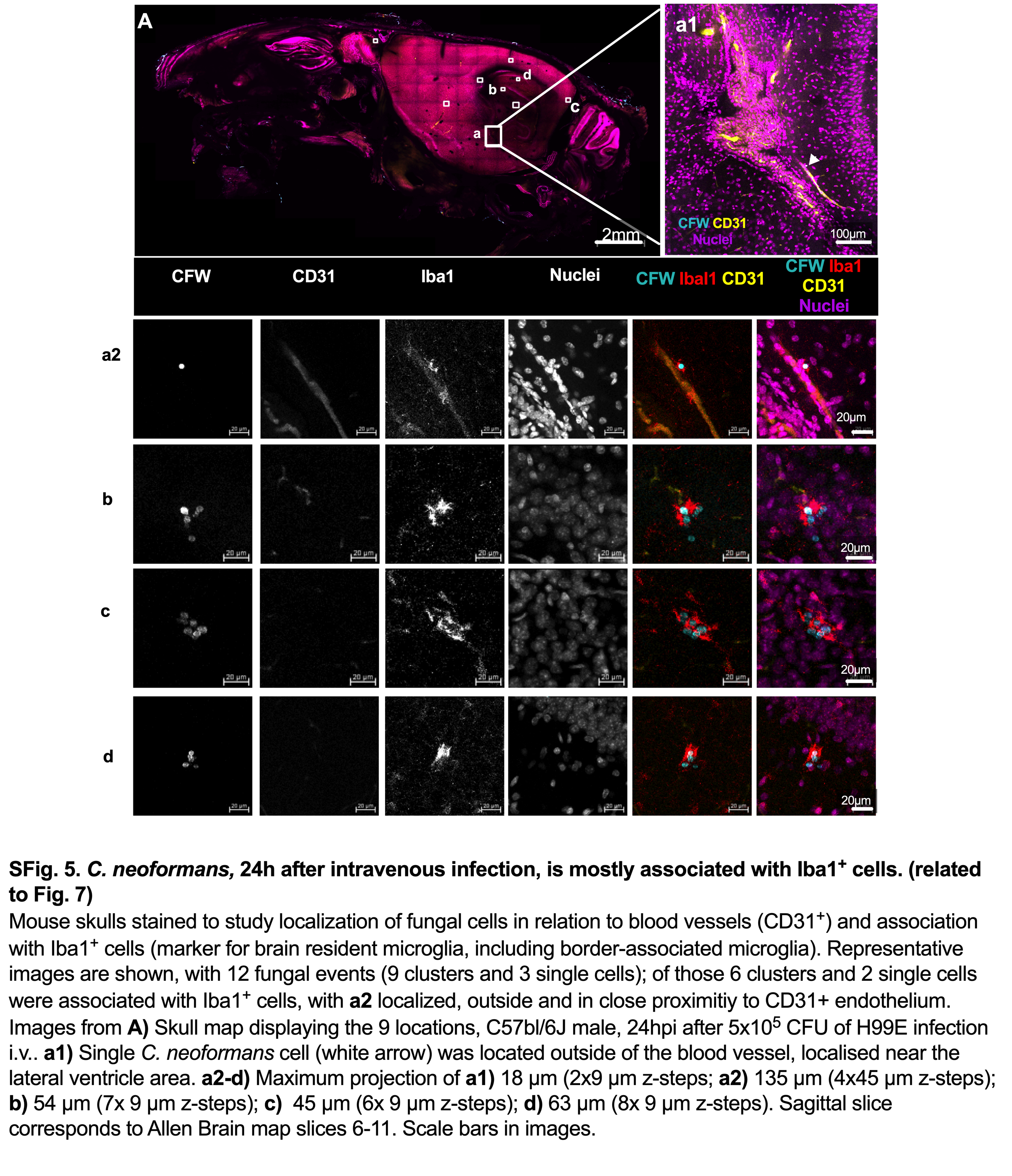

### SFig6. CX3Cr1 and Iba1 coloc.tiff

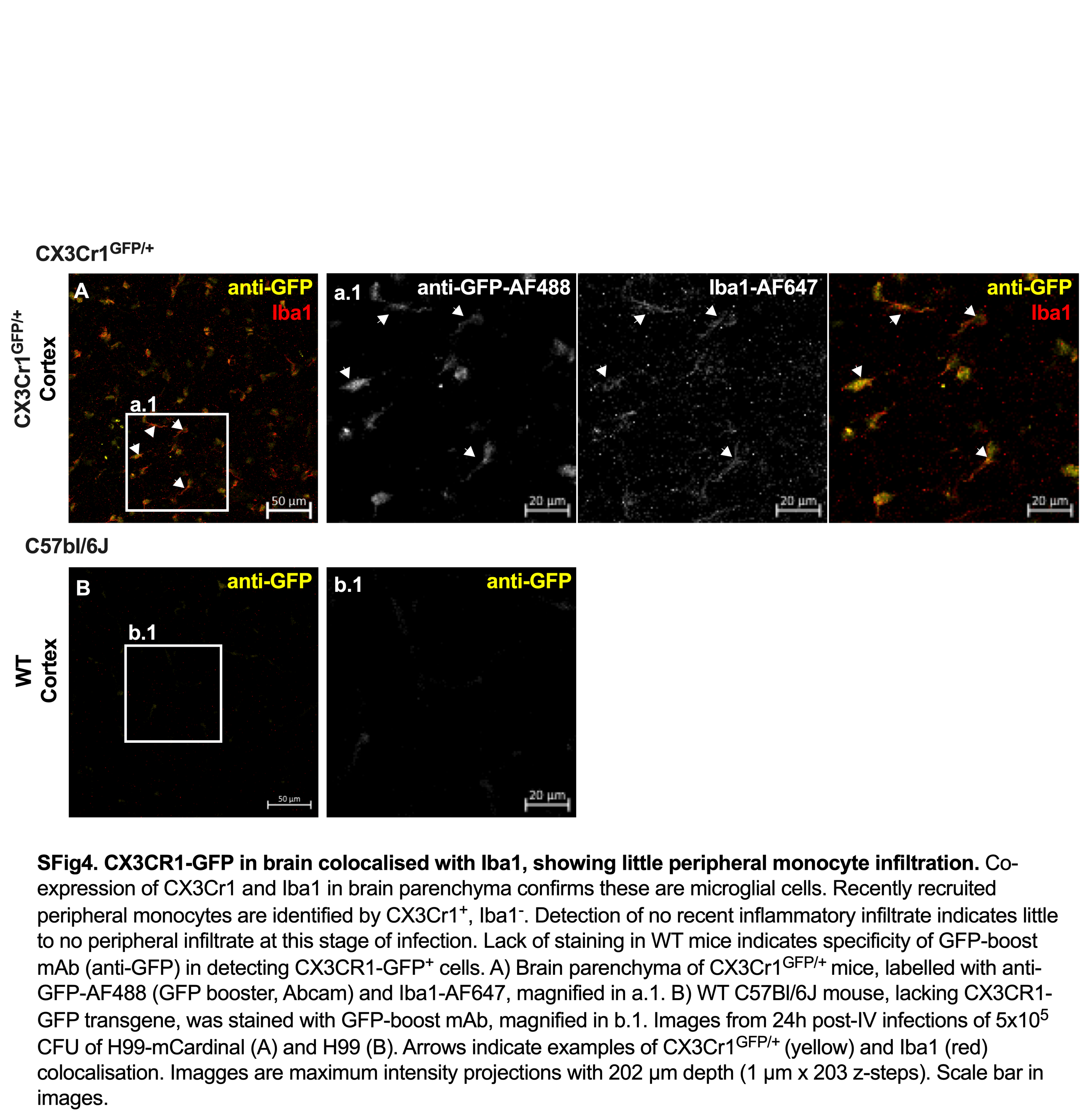

### Sfig7 microglia fused example b1.tiff

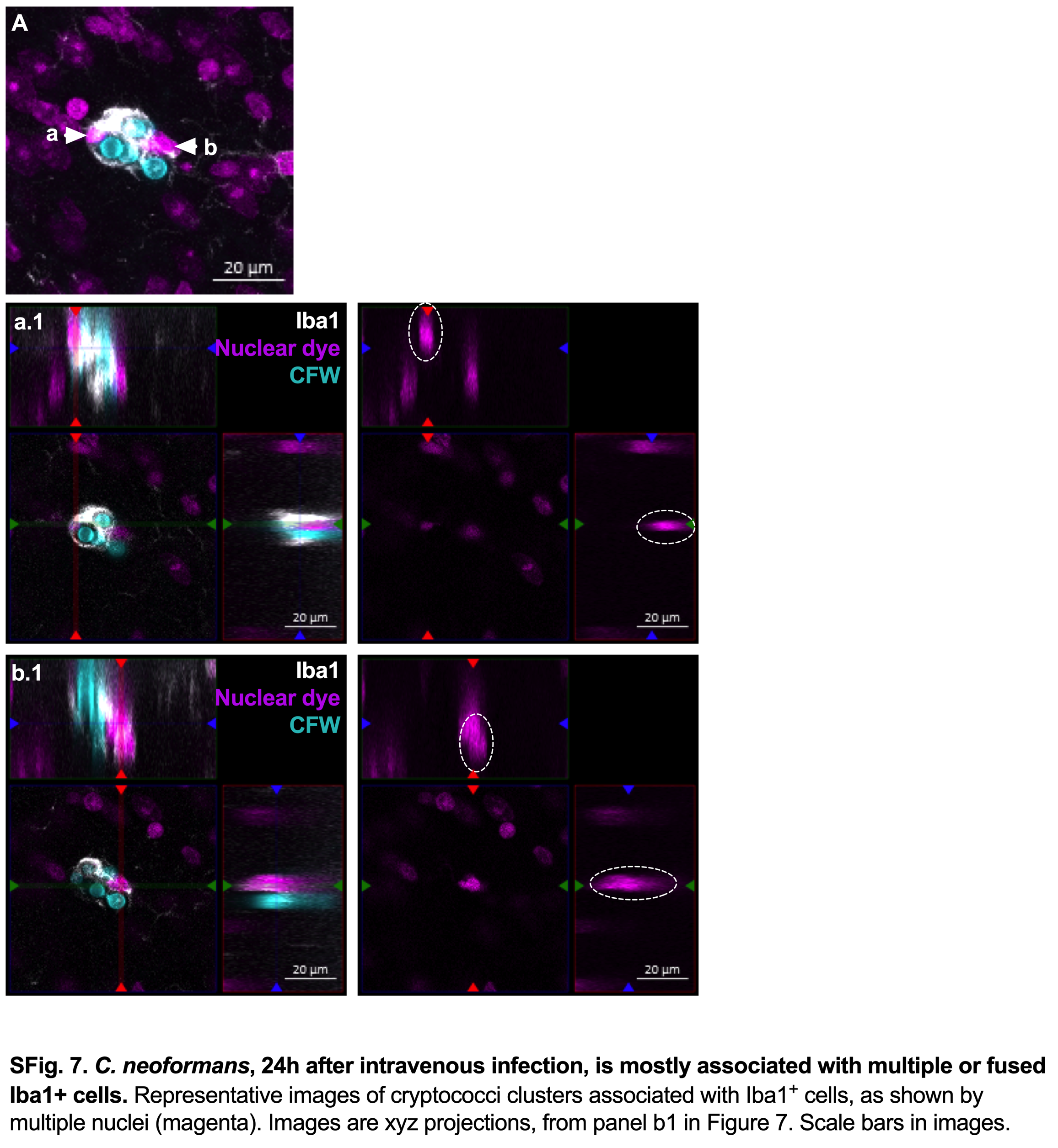

### SFig8. microglia fused example c1.tiff

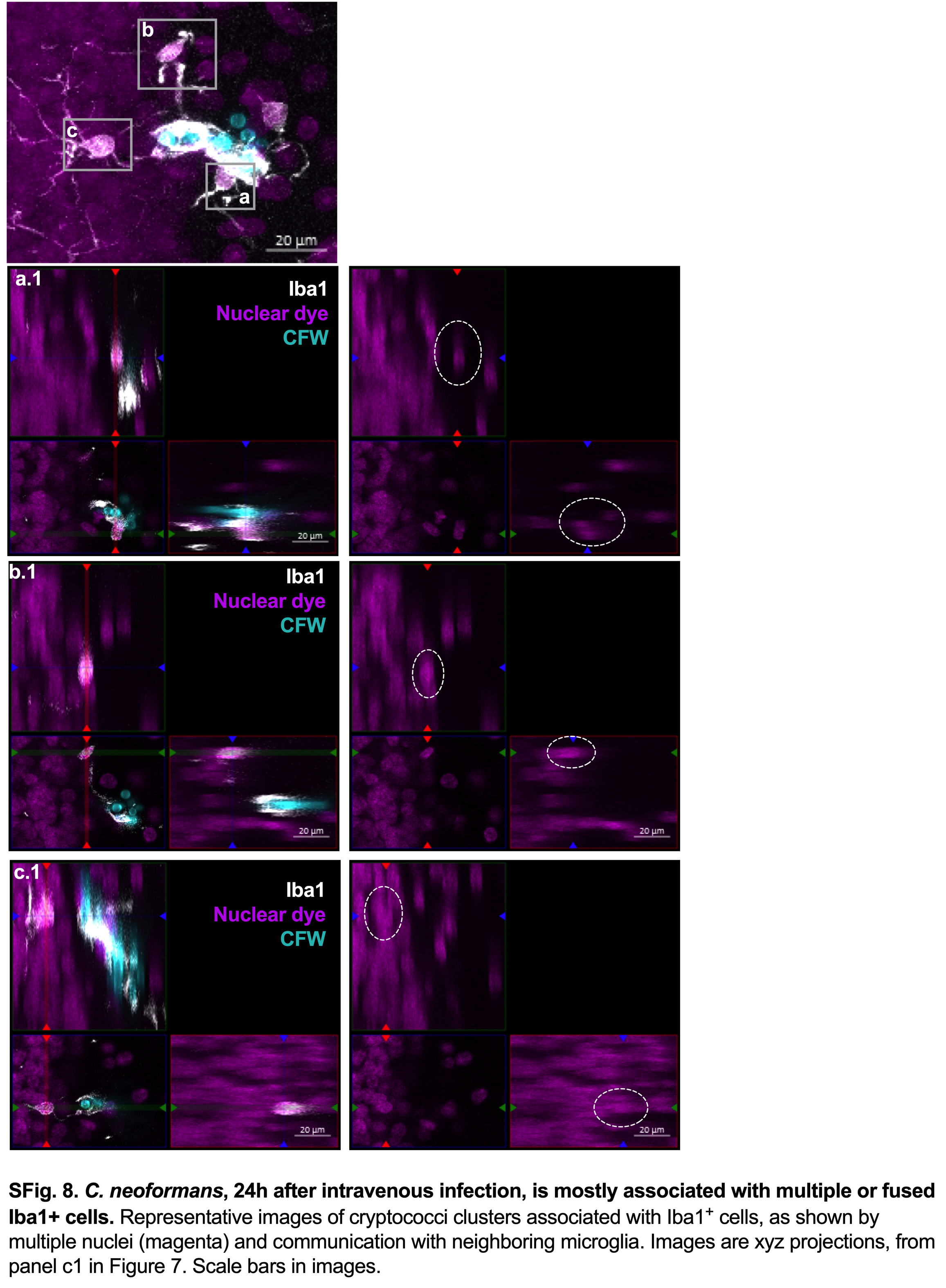
